# Link Prediction through Deep Generative Model

**DOI:** 10.1101/247577

**Authors:** Xu-Wen Wang, Yize Chen, Yang-Yu Liu

## Abstract

Inferring missing links or predicting future ones based on the currently observed network is known as link prediction, which has tremendous real-world applications in biomedicine^1–3^, e-commerce^4^, social media^5^ and criminal intelligence^6^. Numerous methods have been proposed to solve the link prediction problem^7–9^. Yet, many of these existing methods are designed for undirected networks only. Moreover, most methods are based on domain-specific heuristics^10^, and hence their performances differ greatly for networks from different domains. Here we developed a new link prediction method based on deep generative models^11^ in machine learning. This method does not rely on any domain-specific heuristic and works for general undirected or directed complex networks. Our key idea is to represent the adjacency matrix of a network as an image and then learn hierarchical feature representations of the image by training a deep generative model. Those features correspond to structural patterns in the network at different scales, from small subgraphs to mesoscopic communities^12^. Conceptually, taking into account structural patterns at different scales all together should outperform any domain-specific heuristics that typically focus on structural patterns at a particular scale. Indeed, when applied to various real-world networks from different domains^13–17^, our method shows overall superior performance against existing methods. Moreover, it can be easily parallelized by splitting a large network into several small subnetworks and then perform link prediction for each subnetwork in parallel. Our results imply that deep learning techniques can be effectively applied to complex networks and solve the classical link prediction problem with robust and superior performance.

**Summary:** We propose a new link prediction method based on deep generative models.

---

Networks have become an invaluable tool for describing the architecture of various complex systems, be they of technological, biological, or social in nature^18–21^. Mathematically, any real-world network can be represented by a graph 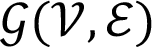, where 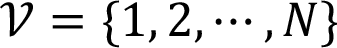 is the node set and 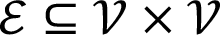 is the link set. A link, denoted as a node pair (*i*, *j*) with 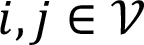, represents certain interaction, association or physical connection between nodes *i* and *j*, which could be either directed or undirected, weighted or unweighted. For many systems (especially biological systems), the discovery and validation of links require significant experimental efforts. Consequently, many real-world networks mapped so-far are substantially incomplete^22,23^. For example, a recent estimate indicates that in human cells the explored protein-protein interactions cover less than 20% of all potential protein-protein interactions^24^. *How to tease out the missing interactions based on the discovered ones?* Moreover, many systems (especially social systems) are very dynamic, as new links are added to the network over time. *How to predict the likelihood of a future interaction between two currently unconnected nodes based on the current snapshot of the network?* Both problems are commonly known as the link prediction problem^25–27^.

An accurate link prediction method will greatly reduce the experimental efforts required to establish the network’s topology and/or accelerate mutually beneficial interactions that would have taken much longer to form serendipitously. Consequently, link prediction has many real-world applications^28,29^. In biomedicine, link prediction can be used to infer protein-protein interactions or drug-target interactions^2^. In e-commerce, it can help build better recommender systems, e.g., Amazon’s “people who bought this also bought” feature^4^. On social media, it can help build potential connections such as the “people you may know” feature on Facebook and LinkedIn^5^. In criminal intelligence analysis, link prediction can assist in identifying hidden co-participation in illicit activities^6^.

Numerous methods, such as similarity-based algorithms^30–34^, maximum likelihood algorithms^7^, probabilistic models^35,36^ have been developed to solve the link prediction problem (see SI Sec.1 for brief descriptions of those existing methods). Yet, many of these existing methods are designed for undirected networks. Moreover, most of these methods are based on domain-specific heuristics, and hence their performances differ greatly for networks from different domains. Here, to quantify the performance of any link prediction method, the standard AUC statistic, i.e., the area under the receiver operating characteristic curve^7,25^, is often employed. To calculate the AUC, we first randomly split the link set *ε* into two parts: (i) a fraction *f* of links as the test or probe set *ε*^*P*^, which will be removed from the network; and (ii) the remaining fraction (1 − *f*) of links as the training set *ε*^*T*^, which will be used to recover the removed links. The AUC statistic is defined to be the probability that a randomly chosen link in the probe set *ε*^*P*^ is given a higher score by the link prediction method than that of a randomly chosen nonexistent link (see SI Sec.2 for details).

A powerful link prediction method that does not rely on any domain-specific heuristic and works for general complex networks has been lacking^37^. Here, we fill the gap by developing a deep learning based link prediction method. Our key idea is to treat the adjacency matrix of a network as the pixel matrix of a binary image. In other words, present (or absent) links will be treated as pixels of value 0 (or 1), respectively. By perturbing the original input network (image) in many different ways through randomly removing a small fraction of present links, we obtain a pool of perturbed networks (images). Those perturbed images will be fed into a deep generative model (DGM) to create fake images that look similar to the input ones (see SI Sec.3 for details of DGMs). Those fake images (networks) will be used to perform link prediction in the original image (network). For the DGM, here we leverage one of the most popular ones --- Generative Adversarial Networks (GANs) that consist of two deep artificial neural networks (called *generator* and *discriminator*) contesting with each other in a game theory framework^11,38^. The generator takes random noise from a known distribution as input and transforms them into fake images through a deconvolutional neural network. The discriminator is a binary classifier (based on a convolutional neural network), which determines whether a given image looks like a real one from the input dataset or like a fake one artificially created by the generator. Over the course of training iterations, the discriminator learns to tell real images from fake ones. At the same time, the generator uses feedback from the discriminator to learn how to produce convincing fake images to fool the discriminator so that it can’t distinguish from real ones (see SI Sec.3 for details). The dissimilarity between fake and real images can be quantified by the so-called Wasserstein distance. During the training process, the generator learns to assign link probabilities to unobserved links (including both missing and nonexistent links) to minimize the Wasserstein distance. (This process is also known as the smooth process^39^). The probabilities assigned to those unobserved links are quite close to zero, but not exactly zero. Hence the generated fake images are grayscale, even though all the input images fed to GAN are binary. If the probabilities assigned to missing links are much higher than that of nonexistent links, then the link prediction is much better than random guess.

To demonstrate our DGM-based link prediction, let’s consider a toy example: a small directed network of 28 nodes and 118 links. Those nodes are labeled appropriately so that the adjacency matrix of this network looks like a binary image of letter **E** with 12 missing pixels, corresponding to 12 missing links (dashed lines in the network, Fig.1). First, we create *M* perturbed binary images by randomly removing a fraction *q* of pixels of value 0 (i.e., those present links) from the original image (network). Second, we use the *M* perturbed binary images as input to train GANs, which will eventually generate *S* fake grayscale images that look similar to the input ones. In this example, we choose *M* = 5,000, *q* = 0.1, and *S* = 500. The existent likelihood of the link between nodes *i* and *j*, denoted as *α*_*ij*_, in the corresponding fake network is simply given by *α*_*ij*_ = 1 − *P*_*ij*_, where *P*_*ij*_ is the rescaled pixel value (ranging from 0 to 1) in each fake grayscale image. Finally, we take the average value 〈*α*_*ij*_〉 = 1 − 〈*P*_*ij*_〉 over all the *S* fake images to get the overall existent likelihood of the link (*i*, *j*). Note that in this toy example all the 12 missing links display higher *α*_*ij*_ than that of nonexistent links, so they are all successfully recovered.

**Fig.1:**
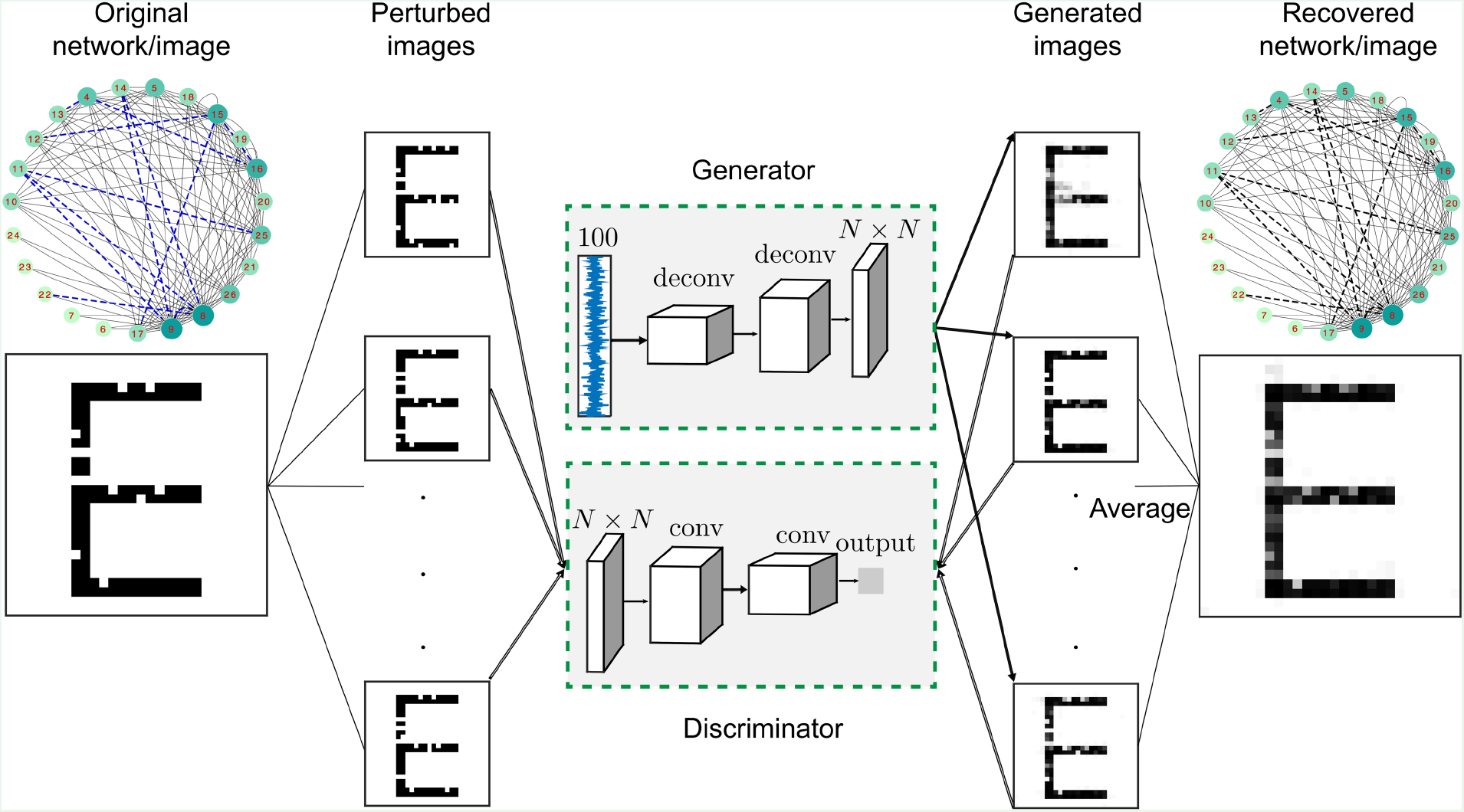
Demonstration of our link prediction method on a directed network. The adjacency matrix of this directed network (with 28 nodes and 118 links) looks like the binary image of letter **E** with 12 missing pixels. Note that 5 isolated nodes are not shown in the network presentation. We perturb the original network (image) by removing 5 links at random in *M* different ways to obtain a pool of perturbed networks (images) *I*_*i*_ (*i* = 1, …, *M*) (*M* = 5000 for this example). This input dataset will be fed into the generative adversarial networks (GANs) that consist of two deep artificial neural networks: generator and discriminator. The generator takes the noise drawn from a uniform distribution (with 100 dimensions for this example) as input and produces fake images. The discriminator is a binary classifier that tells whether a given image is a real one from the input dataset or a fake one produced by the generator. Over the course of training iterations, the generator can produce convincing fake images *P* from the feedback offered by the discriminator. The pixel value *P*_*ij*_ in the fake grayscale image *P* can be used to calculate the existent probability of a link between a node pair: *α*_*ij*_ = 1 − *P*_*ij*_. The final existent probability is calculated by averaging *α*_*ij*_ over *S* (*S* = 500) generated fake networks.

Fig.1 may remind us the classical image inpainting problem, where we need to reconstitute or retouch the missing or damaged regions of an image to make it more legible and to restore its unity^40^. We emphasize that the link prediction problem addressed here is fundamentally different from the image inpainting problem. For image inpainting, we generally know the locations of the damaged regions of an image. While for link prediction, we don’t know which links are missing in a network. In fact, teasing them out is exactly the task of link prediction.

At the first glance, our DGM-based link prediction seems to heavily rely on the existing patterns in the adjacency matrix of the original network. After all, we are treating a network as an image. But do we have to sophisticatedly label the nodes in the network to ensure the success of our method? To address this concern, we perform the following numerical experiment. We start from a network with an appropriate node labeling such that the adjacency matrix looks exactly as the binary image of letter **E** without any missing pixels. (See SI Fig. S1 for a more complicated synthetic network generated by the stochastic block model.) Then we relabel *η* fraction of the nodes in the network so that the binary image associated with its adjacency matrix looks much more random than the letter **E**. Note that the network structure is fixed, and we just label the nodes differently so that the resulting adjacency matrices (or binary images) look quite different. We then compare the performance of our method at different *η* values, as well as the performance of two classical link prediction methods for directed networks that do not depend on the node labeling at all. We find that for this small directed network the performance of our method degrades only slightly even after we relabel 25% nodes (Fig.2a). When we relabel more nodes, the performance is actually quite stable. Even if we relabel all the nodes, the AUC of our method is still about 0.9, which is higher than that of other link prediction methods for directed networks, such as the preferential attachment (PA)^30^ based method (with AUC~0.85) and the low-rank matrix completion (LRMC)^41^ method (with AUC~0.7). In SI Fig.S2, we further show that the AUC of our method is generally above 0.9 with different completely random node labelings of this network. This is simply because that even after completely random node labeling, there are still some small-scale patterns (e.g., many short line segments in the relabeled image of **E**) that can be leveraged by our method.

**Fig.2:**
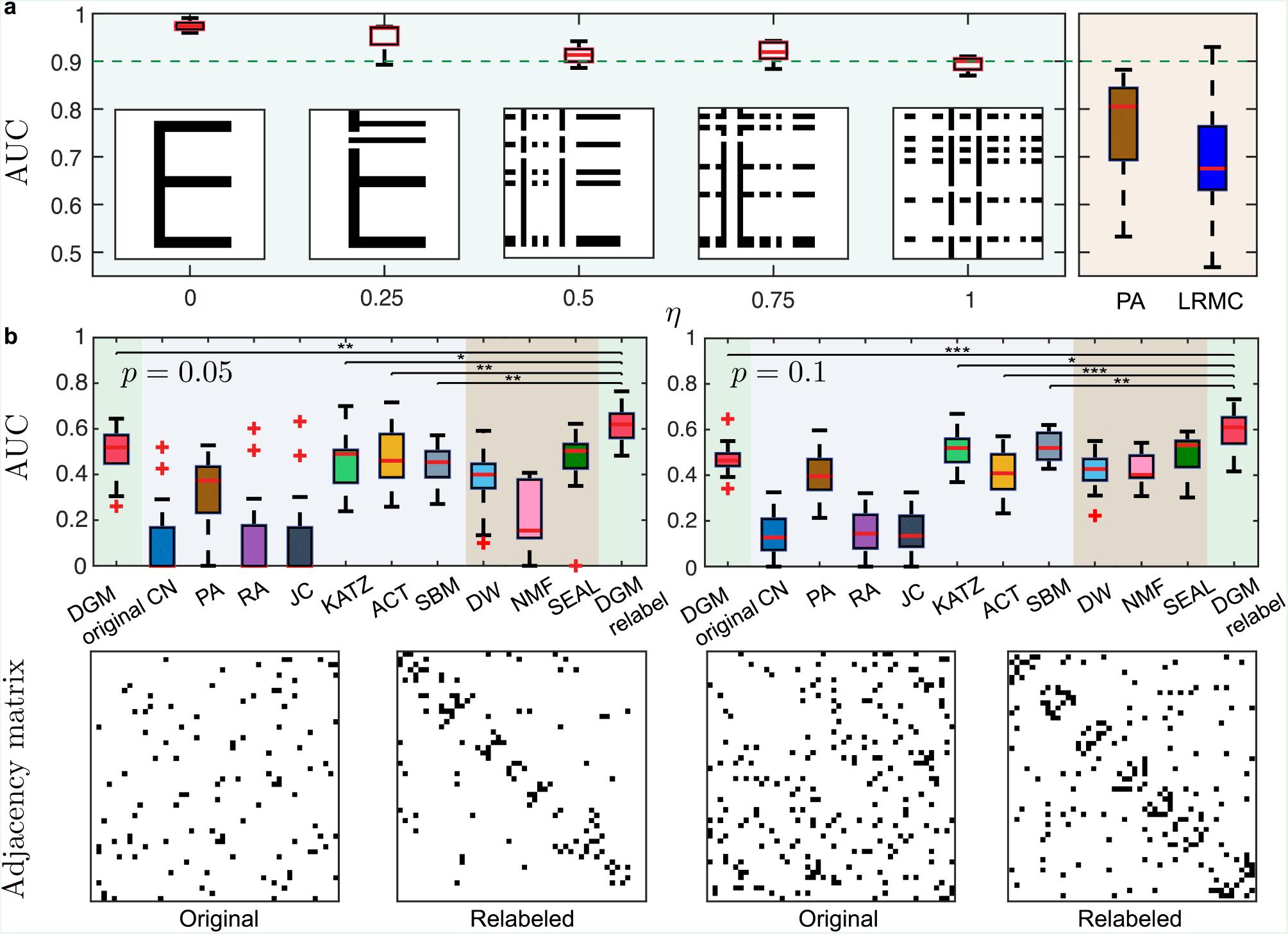
Impact of node labeling on the performance of our DGM-based link prediction method. **a**, A randomly selected fraction of *η* nodes are relabeled in a directed network whose original adjacency matrix looks exactly as the binary image of letter **E**. We randomly divide the links into two parts: a fraction of 10% links chosen as the probe set and the remaining 90% links as the training set. We perform link prediction using three different methods: DGM (deep generative model based), PA (preferential attachment based), and LRMC (low rank matrix completion). In this example, we choose *M* = 1000 for our DGM-based method. Even after we relabel all the nodes so that the adjacency matrix doesn’t display prominent features, the median AUC of our DGM-based method is still around 0.9, while it is 0.85 for the PA method and 0.7 for the LRMC method. **Inset:** The adjacent matrices corresponding to different relabeling fractions, where black pixels represent existing links. **b**, AUCs of DGM-based and other traditional methods in the link prediction of Erdős– Rényi (ER) random graphs (*N* = 48) with different connection probability *p*. The adjacent matrices (before and after node relabeling) at different connection probabilities are also shown. DGM: deep generative model; CN: common neighbors; PA: preferential attachment; RA: resource allocation; JC: Jaccard index; KATZ: Katz index; ACT: average commute time; SBM: stochastic block model; DW: deepwalk embedding method; NMF: non-negative matrix factorization; SEAL: learning from Subgraphs, Embeddings, and Attributes for Link prediction. Asterisks in panel (b) shows whether the AUC of our DGM-based link prediction method is significantly higher than that of the other three traditional algorithms (paired-sample *t*-test). Significance levels: p-value<0.05(*), <0.01(**), <0.001(***).

The results presented in Fig.2a indicate that for those networks that have strong structural patterns, our DGM-based link prediction does not heavily rely on the detailed node labeling. However, to optimize the performance of our method, one should still label the nodes accordingly. This can be achieved by extracting community structure in the network^42,43^, for example, using the classical Louvain method^44^. To test this simple idea, we consider the worst scenario --- random graphs generated from the classical Erdős–Rényi (ER) model, where any two of *N* nodes are randomly connected with probability *p*^45^. By definition, in the large *N* limit, ER random graphs do not display any structural patterns and hence all link prediction methods are doomed to failure. For small *N*, a graph generated by the ER model might display certain structural patterns, but link prediction should still be very challenging, if not impossible. We apply our method as well as various traditional methods to ER random graphs (*N* = 48) at different connection probability *p* and with random node labeling. We find that no link prediction method performs significantly better than random guess, especially for networks with lower connection probability (Fig.2b). However, applying the Louvain method first will capture some structural patterns in the network (emerging as diagonal band structure in the adjacency matrices), which will significantly improve the AUC of our method (Fig. 2b; paired-sample *t*-test). This result suggests that any structural patterns should be exploited for our DGM-based link prediction.

Real-world complex networks certainly display more prominent structural patterns than ER random graphs. Thanks to the deep neural networks in the DGM, our method can actually leverage structural patterns in a real network at different scales all together, from small subgraphs to community structure. To demonstrate this, we consider the character co-occurrence network of Victor Hugo’s *Les Misérables*. As shown in Fig.3a, this network displays many interesting structural patterns, e.g., stars, cliques, and communities. After node labeling using the Louvain method^44^, those structural patterns naturally emerge in the matrix (image) presentation. In particular, those stars show up as line segments, cliques and communities appear as blocks in the image (Fig.3b). After training, deep neural networks with many layers are able to extract the most important structural patterns of the network as the key features of the corresponding image. Note that at the same layer of the deep neural network, different filters can actually learn different feature representations: some focus more on lower level features such as line segments, while others focus more on higher level features such as blocks (Fig.3c). Deeper layers will typically capture higher level features or more global structural patterns (Fig.3d). Leveraging those features at different levels or structural patterns learned at different scales, our DGM-based link prediction performs very well (see SI Sec.4.2.2 and 4.2.3 for more details). Indeed, for this particular network, with a fraction *f* = 0.1 of links as test set, we have AUC~0.95, much higher than that of other link prediction methods, e.g., CN (with AUC~0.7) and SEAL (with AUC~0.85).

**Fig.3:**
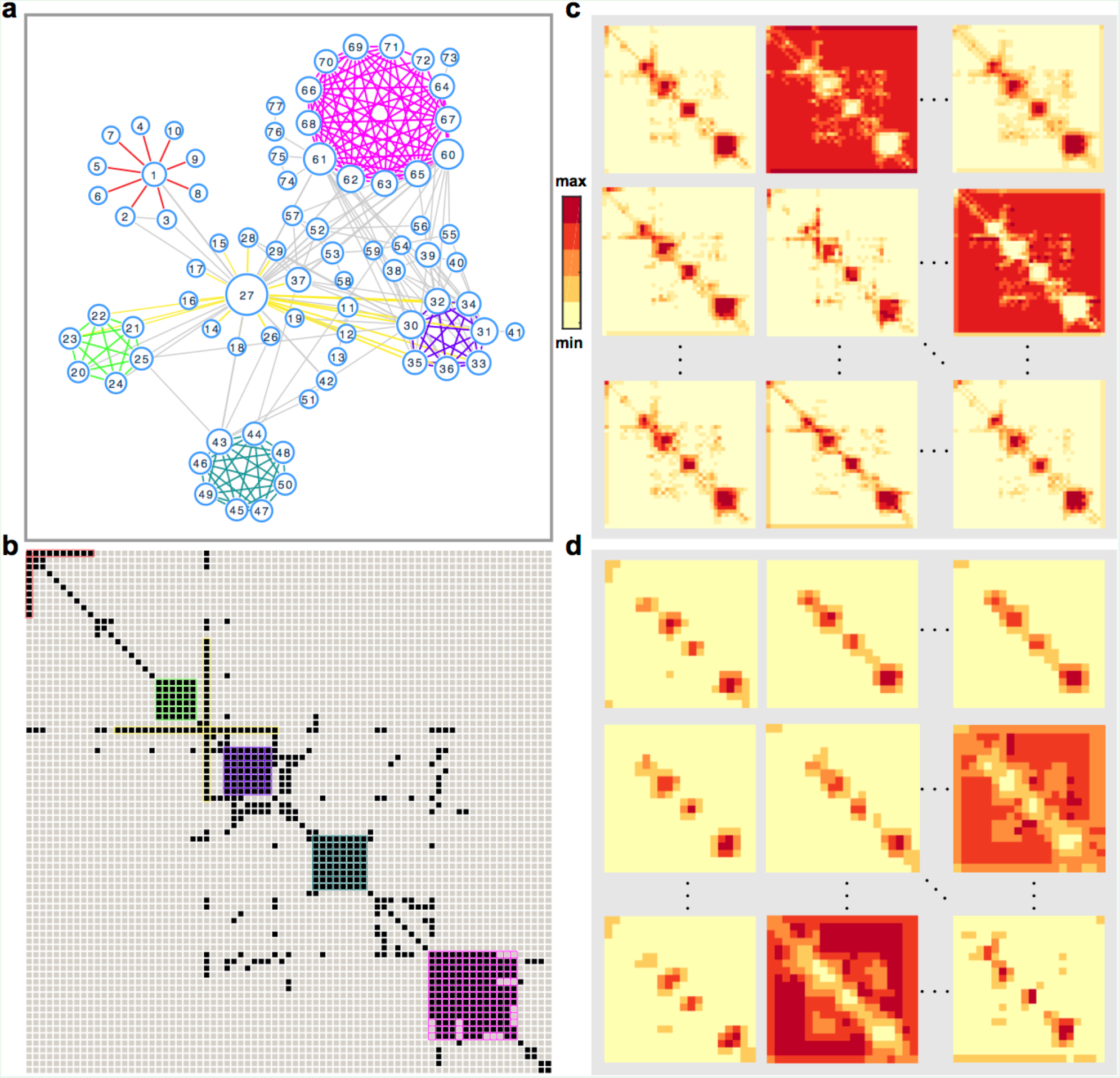
Deep neural networks in the DGM are able to learn different structural patterns of a network at different scales. **a**, The character co-occurrence network of Victor Hugo’s *Les Misérables* (with 77 nodes) contains several interesting structural patterns such as stars, cliques, and community structure. **b**, The matrix (image) representation of the network, with node labeling based on the Louvain method. Those structural patterns are highlighted in different colors. **c**, Learned feature maps from the first convolutional layer with trained filters. There are in total 64 feature maps. Each of them is of size 40 × 40. **d**, Learned feature maps from the second convolutional layer with trained filters. There are in total 128 feature maps. Each of them is of size 20 × 20. For each feature map, higher (or lower) values are shown in redder (or yellower) color.

To systematically demonstrate the advantage of our DGM-based link prediction in real-world applications, we compare the performance of our method with that of both classical and state-of-the-art link prediction methods for a wide range of real-world networks, from social, economic, technological to biological networks (see SI Sec.6 for brief descriptions of real networks analyzed in this work). For undirected networks (Fig.4a), we find that generally global similarity indices (e.g., Katz, ACT) and SBM-based link prediction methods perform better than local similarity indices (e.g., CN, PA, RA) based methods. But the performances of those heuristics-based methods vary a lot over different network domains. Some of them actually perform even worse than random guess, especially when the training set is small (corresponding to large *f*). By contrast, our DGM-based method displays very robust and high performance for various undirected networks. It also outperforms several state-of-the-art link prediction methods based on non-negative matrix factorization^33^, network embedding^34^, and graph neural networks^46^. For directed networks (Fig.4b), most of the existing methods (especially those state-of-the-art methods) are actually not applicable, except two classical methods: PA and LRMC. We compare the performance of our method with those of PA and LRMC. Again, we find that our method displays more robust and better performance than PA and LRMC for various directed networks.

**Fig.4:**
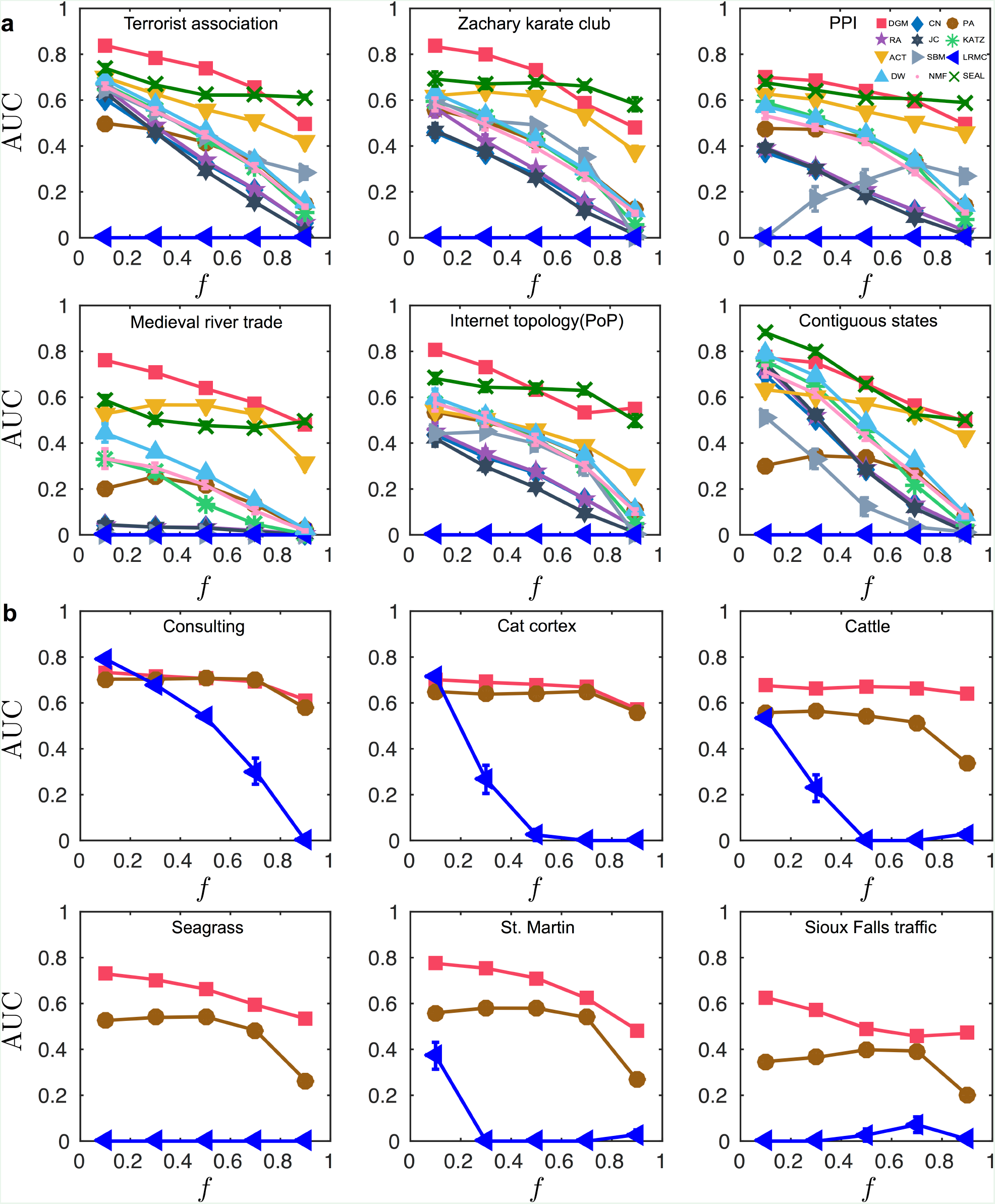
Our DGM-based link prediction displays very robust and high performance for both undirected and directed real-world networks. DGM: deep generative model based link prediction; CN: common neighbors; PA: preferential attachment; RA: resource allocation; JC: Jaccard index; KATZ: Katz index; ACT: average commute time; SBM: stochastic block model; LRMC: low rank matrix completion; DW: deepwalk embedding method; NMF: non-negative matrix factorization; SEAL: learning from Subgraphs, Embeddings, and Attributes for Link prediction (see SI Sec.1 for details of each algorithm). **a**, Undirected networks. Top: Terrorist association network, Zachary karate club, Protein-protein interaction (PPI) network (a subnetwork of PPIs in *S. cerevisiae*). Bottom: Medieval river trade network in Russia, Internet topology (at the PoP level), Contiguous states in USA. **b**, Directed networks. Top: Consulting (a social network of a consulting company), cat cortex (the connection network of cat cortical areas), cattle (a network of dominance behaviors of cattle). Bottom: Seagrass food web, St. Martin food web, Sioux Falls traffic network. AUC of our DGM-based method is the average AUC over the last 20 epochs of the total 150 epochs for all of networks. Here an epoch is one full training cycle on the training set. For all the undirected real networks, we apply the Louvain method first to label the nodes appropriately. Directed networks are labeled by the method proposed in Ref. [48]. Error bar represents the standard error of the mean (s.e.m.).

We emphasize that before applying our DGM-based link prediction to each of the real-world networks tested in Fig.4, we performed node labeling (also known as matrix reordering in literature^47^) to get the matrix (or image) presentation of the network. For the sake of simplicity, we just labeled nodes in a network based on its community structure. In particular, for undirected networks, we apply the Louvain method; for directed networks, we apply the method proposed in Ref. [48]. We emphasize that our approach doesn’t rely on the presence of communities in a network. Any structural patterns at different scales (from small subgraphs to communities, see Fig.3) can be and should be leveraged all together. A reasonable node labeling can actually be achieved in many different ways other than just community detection. For example, we developed a simulated annealing based node labeling method to maximize the “compactness” of the image (see SI Sec.5.3). We found that for real-world networks the performance of our DGM-based link prediction actually does not heavily depend on the specific node labeling algorithm (see SI Sec.5 and Figs.S11, S12, S13). This is consistent with the results presented in Fig.2a, where we show that as long as the network has strong structural patterns, then any reasonable node labeling will offer a plausible matrix (or image) presentation of the network, which can be used for our DGM-based link prediction.

Since our method essentially treats a network as an image, it can be easily parallelized by splitting a large network (image) into several small subnetworks (subimages) and then performing link prediction for each subnetwork (subimage) in parallel (Fig.5a). Note that node labeling is the first step of our approach. In this step, we always treat the whole network as an image (by any reasonable node labeling algorithm), regardless of the network size. Training the deep generative model (DGM) is the second step of our approach. In this step, if the network/image is small, we train the DGM and hence perform link prediction for the whole image. Only if the network/image is too large, for which the DGM cannot be easily trained, we need to split the image into subimages, train DGM and perform link prediction for different subimages in parallel. This splitting typically does not decrease the overall link prediction performance, compared with the result of treating the large network as a whole (Fig. 5b). For each subnetwork, when only the information of the subnetwork is provided, our method outperforms other methods (Fig.5b). In fact, even if other methods (e.g., PA and LRMC) use the information of the whole network to perform link prediction for a subnetwork, our method that only relies on the information of the subnetwork still display better performance (Fig.5b).

**Fig.5:**
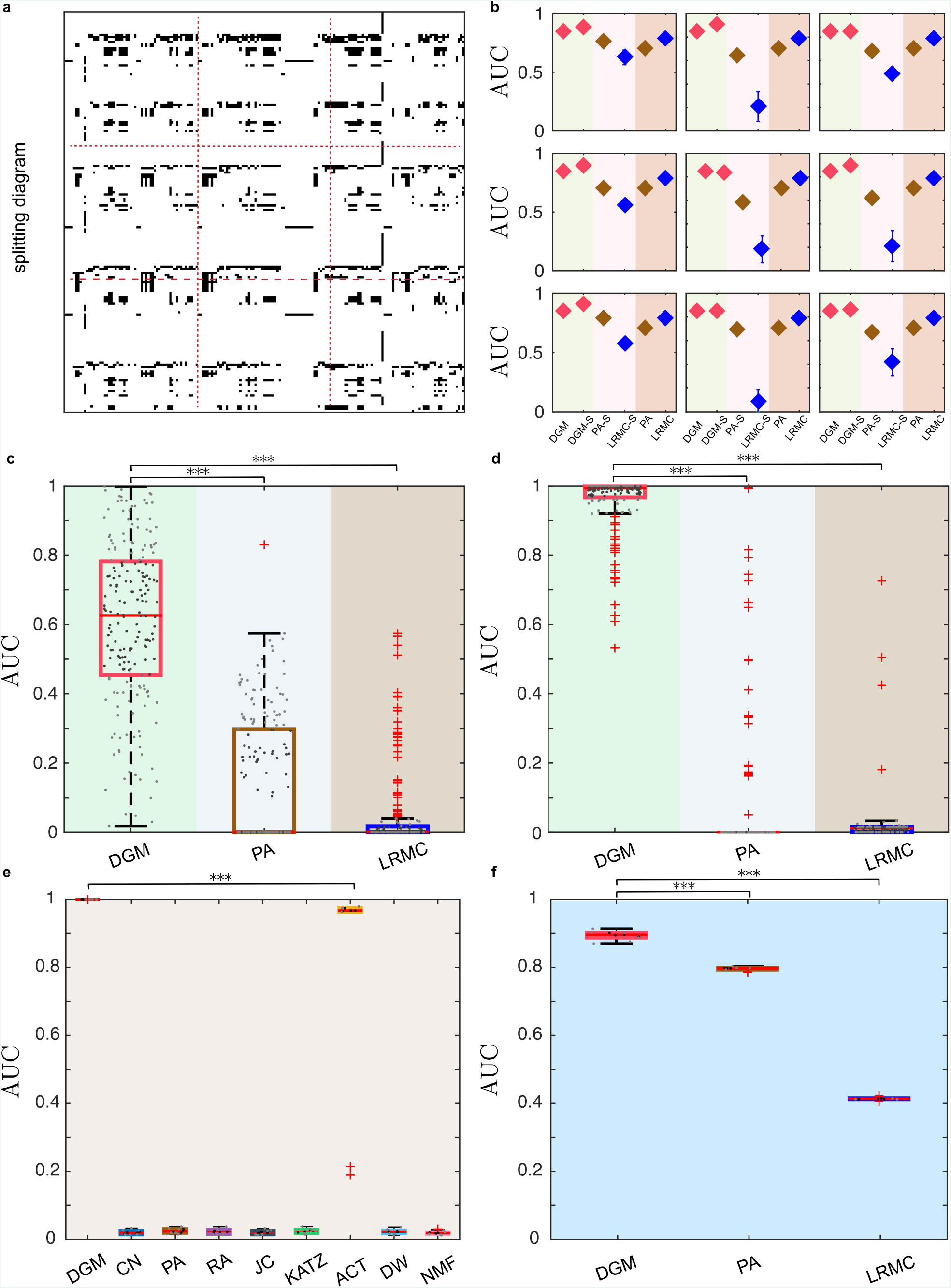
The DGM-based link prediction method can infer missing links of large networks and arbitrarily selected subnetworks within large networks. **a**, The adjacency matrix of a real network: The Little Rock food web. The network (image) is split into 9 subnetworks (subimages). **b**, AUC of DGM, PA, and LRMC on the original network and AUC of DGM, PA, and LRMC on each subnetwork (DGM-S, PA-S, LRMC-S). **c-d**, We perform link prediction for 200 randomly selected subnetworks (of size 60) chosen from two large-scale real networks: (e) Facebook wall posts (with 46,952 nodes and 87,6993 links) and (f) Google+ (with 23,628 nodes and 39,242 links), respectively. We randomly divide the links of the relabeled networks into two parts: a fraction of 10% links are chosen as probe set and the remaining 90% fraction of links as training set (here, each subnetwork contains 15 links at least). Asterisks at the top of each panel shows whether the AUC of our DGM-based link prediction model is significantly higher than that of the other two traditional algorithms (paired-sample *t*-test). Significance levels: p-value<0.05(*), <0.01(**), <0.001(***), not significant (NS). **e-f**, AUC of DGM and other scalable link prediction methods in two large-scale networks: (e) Facebook-NIPS (with *N* = 2,888 nodes and 2,981 links), and (f) US airports (with *N* = 1,574 nodes and 28,236 links).

The image representation of complex networks also allows us to focus on any specific subnetwork of interest and just predict the missing links in that subnetwork. For example, we perform link prediction for 200 subnetworks of size 60 randomly selected from two large real networks: Facebook wall posts^49^ (with 46,952 nodes and 87,6993 links) and Google+^50^ (with 23,628 nodes and 39,242 links). We find that our method shows much higher AUC than other methods (Fig. 5c,d). All these results suggest that our method holds great promise in link prediction for large-scale real-world networks.

To directly demonstrate the performance of our method in analyzing large-scale networks, we consider an undirected network: Facebook-NIPS (with *N* = 2,888 nodes), and a directed network: US airports (with *N* = 1,574 nodes). We randomly remove a fraction (*f* = 0.1) of links as test set. To facilitate the training process of DGM and speed up the link prediction, we still split each large network (image) into several small subnetworks (subimages), and then perform link prediction for each subnetwork. But, in the end, to have a fair comparison with other methods (that always treat the large network as a whole), we calculate the AUC of our method from the whole network (constructed by merging subnetworks/subimages generated by the DGM). We find that clearly for both large-scale real networks (Facebook-NIPS and US airports) our method outperforms other existing methods (Fig. 5e,f). See SI Fig.S14 for results of large-scale model networks.

In summary, our DGM-based link prediction shows superior performance against existing methods for various types of networks, be they of technological, biological, or social in nature. Since our method treats the adjacency matrix of a network as an image, it can be naturally extended to solve the link prediction problem for bipartite graphs, multi-layer networks and multiplex networks, where the adjacency matrices have certain inherent structure. With small modification, it can also be used to perform link prediction in weighted graphs (see SI Fig.S15). In principle, any DGM can be utilized in our method. But we find that, for the link prediction purpose, GANs perform much better than other DGMs, e.g., variational autoencoder^51^ (see SI Fig.S16). There are several hyperparameters in training the GANs (see SI Sec.4 for details). In this work, we use the same set of hyperparameters for all the networks to show a conservative AUC estimation of our method. The performance of our method can certainly be further improved by carefully tuning those hyperparameters for a specific network of interest.

We should admit that although our DGM-based link prediction displays superior performance in various real-world networks, its time complexity is higher than traditional heuristic-based methods (e.g., Common Neighbors^52^, Preferential Attachment^30^) and embedding-based methods (e.g., DeepWalk^34^, node2vec^53^). (See SI Sec.4.2.4 for detailed analysis of its time complexity.) Such a *speed-accuracy tradeoff* deserves a very careful consideration in real-world applications. For certain link prediction applications, such as recommendation system in e-commerce or online social media with daunting network sizes, speed is the major concern, hence traditional link prediction methods still have big advantages. For applications in biomedicine (e.g., inferring protein-protein interactions or drug-target interactions) or criminal intelligence analysis (e.g., identifying hidden accomplice in criminal activity), those networks are much smaller than social media networks, and accuracy is way more important than speed. In those cases, we anticipate that our DGM-based link prediction should have an unparalleled advantage. Furthermore, we suggest that one should definitely exploit graphics process unit parallelism to train the GANs^54^, which will certainly speed up our method. Finally, we emphasize that, in real-world applications of link prediction, any additional side-information, such as node attributes, can be incorporated into our method to further improve the link prediction.

Recently, the notion of permutation-invariance has been discussed a lot in the deep learning literature^55–59^. A network method is called permutation-invariant if it produces the same output regardless of the node labeling used to encode the adjacency matrix of the network. We should admit that, since we treat the adjacency matrix of a network as a binary image, our method is by definition not permutation invariant. But we emphasize that our method is approximately permutation invariant in practice. As shown in Figs.S12, S13, node labeling doesn’t significantly affect the performance of our link prediction method, as long as the node labeling method leverages existing structure features in the network.

## Acknowledgments

We thank Yizhou Sun, Tina Eliassi-Rad, Pan Zhang, Changjun Fan, Nima Dehmamy, Santo Fortunato, Yong-Yeol Ahn, Huawei Shen and Marco Tulio Angulo for valuable discussions.

## Contributions

Y.-Y.L. conceived and designed the project. X.-W.W. and Y.C. did the analytical and numerical calculations. X.-W.W. analyzed all the real networks. All authors analyzed the results. Y.-Y.L. and X.-W.W. wrote the manuscript. Y.C. edited the manuscript.

## Author Information

The authors declare no competing financial interests. Correspondence and requests for materials should be addressed to Y.-Y.L.

## References

1. Campillos, M., Kuhn, M., Gavin, A.-C., Jensen, L. J. & Bork, P. Drug Target Identification Using Side-Effect Similarity. Science 321, 263–266 (2008).

2. Luo, Y. et al. A network integration approach for drug-target interaction prediction and computational drug repositioning from heterogeneous information. Nat. Commun. 8, (2017).

3. Liu, F., Liu, B., Sun, C., Liu, M. & Wang, X. Deep learning approaches for link prediction in social network servicess. in 425–432 (Springer, 2013).

4. Linden, G., Smith, B. & York, J. Amazon. com recommendations: Item-to-item collaborative filtering. IEEE Internet Comput. 7, 76–80 (2003).

5. Blagus, N., Šubelj, L. & Bajec, M. Self-similar scaling of density in complex real-world networks. Phys. Stat. Mech. Its Appl. 391, 2794–2802 (2012).

6. Berlusconi, G., Calderoni, F., Parolini, N., Verani, M. & Piccardi, C. Link prediction in criminal networks: a tool for criminal intelligence analysis. PloS One 11, e0154244 (2016).

7. Guimerà, R. & Sales-Pardo, M. Missing and spurious interactions and the reconstruction of complex networks. Proc. Natl. Acad. Sci. 106, 22073–22078 (2009).

8. Friedman, N., Getoor, L., Koller, D. & Pfeffer, A. Learning probabilistic relational models. in IJCAI 99, 1300–1309 (1999).

9. Kovács, I. A. et al. Network-based prediction of protein interactions. bioRxiv 275529 (2018).

10. Sarukkai, R. R. Link prediction and path analysis using Markov chains1. Comput. Netw. 33, 377–386 (2000).

11. Goodfellow, I. et al. Generative adversarial nets. in Advances in neural information processing systems 2672–2680 (2014).

12. Girvan, M. & Newman, M. E. Community structure in social and biological networks. Proc. Natl. Acad. Sci. 99, 7821–7826 (2002).

13. Zhang, B., Liu, R., Massey, D. & Zhang, L. Collecting the Internet AS-level topology. ACM SIGCOMM Comput. Commun. Rev. 35, 53–61 (2005).

14. Krebs, V. E. Mapping networks of terrorist cells. Connections 24, 43–52 (2002).

15. Baird, D., Luczkovich, J. & Christian, R. R. Assessment of spatial and temporal variability in ecosystem attributes of the St Marks National Wildlife Refuge, Apalachee Bay, Florida. Estuar. Coast. Shelf Sci. 47, 329–349 (1998).

16. Christian, R. R. & Luczkovich, J. J. Organizing and understanding a winter’s seagrass foodweb network through effective trophic levels. Ecol. Model. 117, 99–124 (1999).

17. LeBlanc, L. J., Morlok, E. K. & Pierskalla, W. P. An efficient approach to solving the road network equilibrium traffic assignment problem. Transp. Res. 9, 309–318 (1975).

18. Albert, R. & Barabási, A.-L. Statistical mechanics of complex networks. Rev. Mod. Phys. 74, 47 (2002).

19. Newman, M. E. The structure and function of complex networks. SIAM Rev. 45, 167–256 (2003).

20. Boccaletti, S., Latora, V., Moreno, Y., Chavez, M. & Hwang, D. Complex networks: Structure and dynamics. Phys. Rep. 424, 175–308 (2006).

21. Scholtes, I. et al. Causality-driven slow-down and speed-up of diffusion in non-Markovian temporal networks.Nat. Commun. 5, 5024 (2014).

22. Von Mering, C. et al. Comparative assessment of large-scale data sets of protein–protein interactions. Nature 417, 399–403 (2002).

23. Han, J.-D. J., Dupuy, D., Bertin, N., Cusick, M. E. & Vidal, M. Effect of sampling on topology predictions of protein-protein interaction networks. Nat. Biotechnol. 23, 839–844 (2005).

24. Sahni, N. et al. Widespread macromolecular interaction perturbations in human genetic disorders. Cell 161, 647–660 (2015).

25. Clauset, A., Moore, C. & Newman, M. E. J. Hierarchical structure and the prediction of missing links in networks. Nature 453, 98–101 (2008).

26. Lü, L., Pan, L., Zhou, T., Zhang, Y.-C. & Stanley, H. E. Toward link predictability of complex networks. Proc. Natl. Acad. Sci. 112, 2325–2330 (2015).

27. Liben-Nowell, D. & Kleinberg, J. The link-prediction problem for social networks. J. Am. Soc. Inf. Sci. Technol. 58, 1019–1031 (2007).

28. Hulovatyy, Y., Solava, R. W. & Milenković, T. Revealing missing parts of the interactome via link prediction. PloS One 9, e90073 (2014).

29. Martínez, V., Berzal, F. & Cubero, J.-C. A survey of link prediction in complex networks. ACM Comput. Surv. CSUR 49, 69 (2016).

30. Barabási, A.-L. & Albert, R. Emergence of scaling in random networks. science 286, 509–512 (1999).

31. Katz, L. A new status index derived from sociometric analysis. Psychometrika 18, 39–43 (1953).

32. Zhou, T., Lü, L. & Zhang, Y.-C. Predicting missing links via local information. Eur. Phys. J. B 71, 623–630 (2009).

33. Chen, B., Li, F., Chen, S., Hu, R. & Chen, L. Link prediction based on non-negative matrix factorization. PLOS ONE 12, e0182968 (2017).

34. Perozzi, B., Al-Rfou, R. & Skiena, S. DeepWalk: Online Learning of Social Representations. ArXiv14036652 Cs 701–710 (2014). doi:10.1145/2623330.2623732

35. Heckerman, D., Meek, C. & Koller, D. Probabilistic entity-relationship models, PRMs, and plate models. Introd. Stat. Relational Learn. 201–238 (2007).

36. Chaney, A. J., Blei, D. M. & Eliassi-Rad, T. A probabilistic model for using social networks in personalized item recommendation. in Proceedings of the 9th ACM Conference on Recommender Systems 43–50 (ACM, 2015).

37. Martínez, V., Berzal, F. & Cubero, J.-C. A Survey of Link Prediction in Complex Networks. ACM Comput. Surv. 49, 1–33 (2016).

38. Arjovsky, M., Chintala, S. & Bottou, L. Wasserstein GAN. ArXiv170107875 Cs Stat (2017).

39. Yeh, R. A. et al. Semantic image inpainting with deep generative models. in Proceedings of the IEEE Conference on Computer Vision and Pattern Recognition 5485–5493 (2017).

40. Bertalmio, M., Sapiro, G., Caselles, V. & Ballester, C. Image inpainting. in Proceedings of the 27th annual conference on Computer graphics and interactive techniques 417–424 (ACM Press/Addison-Wesley Publishing Co., 2000).

41. Pech, R., Hao, D., Pan, L., Cheng, H. & Zhou, T. Link Prediction via Matrix Completion. EPL Europhys. Lett. 117, 38002 (2017).

42. Newman, M. E. & Girvan, M. Finding and evaluating community structure in networks. Phys. Rev. E 69, 026113 (2004).

43. Radicchi, F., Castellano, C., Cecconi, F., Loreto, V. & Parisi, D. Defining and identifying communities in networks. Proc. Natl. Acad. Sci. U. S. A. 101, 2658–2663 (2004).

44. Blondel, V. D., Guillaume, J.-L., Lambiotte, R. & Lefebvre, E. Fast unfolding of communities in large networks. J. Stat. Mech. Theory Exp. 2008, P10008 (2008).

45. ERDdS, P. & R&WI, A. On random graphs I. Publ Math Debr. 6, 290–297 (1959).

46. Zhang, M. & Chen, Y. Link Prediction Based on Graph Neural Networks. ArXiv Prepr. ArXiv180209691 (2018).

47. Behrisch, M., Bach, B., Henry Riche, N., Schreck, T. & Fekete, J.-D. Matrix Reordering Methods for Table and Network Visualization. Comput. Graph. Forum 35, 693–716 (2016).

48. Arenas, A., Fernández, A. & Gómez, S. Analysis of the structure of complex networks at different resolution levels. New J. Phys. 10, 053039 (2008).

49. Viswanath, B., Mislove, A., Cha, M. & Gummadi, K. P. On the evolution of user interaction in facebook. in Proceedings of the 2nd ACM workshop on Online social networks 37–42 (ACM, 2009).

50. Leskovec, J. & Mcauley, J. J. Learning to discover social circles in ego networks. in Advances in neural information processing systems 539–547 (2012).

51. Sohn, K., Lee, H. & Yan, X. Learning structured output representation using deep conditional generative models. in Advances in Neural Information Processing Systems 3483–3491 (2015).

52. Zhou, T. et al. Solving the apparent diversity-accuracy dilemma of recommender systems. Proc. Natl. Acad. Sci. 107, 4511–4515 (2010).

53. Grover, A. & Leskovec, J. node2vec: Scalable Feature Learning for Networks. in Proceedings of the 22nd ACM SIGKDD International Conference on Knowledge Discovery and Data Mining - KDD ’16 855–864 (ACM Press, 2016). doi:10.1145/2939672.2939754

54. Im, D. J., Ma, H., Kim, C. D. & Taylor, G. Generative Adversarial Parallelization. ArXiv Prepr. ArXiv161204021 (2016).

55. Van Laarhoven, T. & Marchiori, E. in Journal of Machine Learning Research Vol. 15 193–215 (2014).

56. Wu, Z. et al. arXiv:1901.00596v1

57. Meltzer, P., Mallea, M. D. G. & Bentley, P. J. arXiv:1905.03046

58. Maron, H., Ben-Hamu, H., Shamir, N. & Lipman, Y. in 7th International Conference on Learning Representations, ICLR 2019 1–14 (2019).

59. Atamna, A., Sokolovska, N., and Crivello, J. hal-02093451

