## Supplementary Information for "Link Prediction through Deep Generative Model" (from another version of this preprint: version 3, 2019-12-26)

### LINK PREDICTION THROUGH DEEP GENERATIVE MODEL
#### SUPPLEMENTARY MATERIALS

XU-WEN WANG[1], YIZE CHEN[2], and YANG-YU LIU[1,3]

Supplementary Text

Figs. S1 to S16

Tables S1 to S2

References *(56-147)*

---

*Date*: December 10, 2019

[1]Channing Division of Network Medicine, Brigham and Women's Hospital, and Harvard Medical School, Boston, MA 02115, USA

[2]Department of Electrical Engineering, University of Washington, Seattle, WA 98195, USA

[3]Center for Cancer Systems Biology, Dana-Farber Cancer Institute, Boston, MA 02115, USA.

5.2 Node relabeling algorithm based on multiple resolution modular detection . . 23

#### 1. EXISTING LINK PREDICTION ALGORITHMS

A real-world network can be mathematically represented by a graph $G(\mathcal{V}, \mathcal{E})$, where $\mathcal{V} = \{1, 2, \ldots, N\}$ is the node set and $\mathcal{E} \subseteq \mathcal{V} \times \mathcal{V}$ is the link set. A link is a node pair $(i, j)$ with $i, j \in \mathcal{V}$, representing certain interaction, association or physical connection between nodes $i$ and $j$. Link prediction aims to infer the missing links or predict future links between currently unconnected nodes based on the observed links [56, 57, 58]. Many algorithms have been developed to solve the link prediction problem [59, 60, 61, 62]. Here we briefly describe some classical link prediction algorithms.

##### 1.1. Similarity-based algorithms.

For similarity-based algorithms, each non-observed node pair $(i, j)$ is assigned a similarity score $s_{ij}$. Higher score is assumed to represent higher link existence probability. The similarity score can be defined in many different ways.

**(1) Common Neighbors**. The common neighbors algorithm quantifies the overlap or similarity of two nodes as follows [63]:

$$s_{ij} = |\Gamma(i) \cap \Gamma(j)|, \tag{S1}$$

where $\Gamma(i)$ denotes the set of neighbors of node $i$, $\cap$ denotes the intersection of two sets and $|X|$ denotes the cardinality or size of set $X$.

**(2) Jaccard Index**. The Jaccard index measures the overlap of two nodes with normalization [64]:

$$s_{ij} = \frac{|\Gamma(i) \cap \Gamma(j)|}{|\Gamma(i) \cup \Gamma(j)|}, \tag{S2}$$

where $\cup$ denotes the union of two sets.

**(3) Preferential Attachment Index**. This index assumes that the existent likelihood of a link between two nodes is proportional to the product of their degrees [65]:

$$s_{ij} = k_i \times k_j, \tag{S3}$$

where $k_i$ is the degree of node $i$.

**(4) Resource Allocation Index**. This index is based on a resource allocation process between pair of nodes [66, 67]. Consider a node pair $(i, j)$, the similarity between $i$ and $j$ is defined as the amount of resource $j$ received from $i$ through their common neighbors:

(S4) $$s_{ij} = \sum_{m \in \Gamma(i) \cap \Gamma(j)} \frac{1}{k_m}.$$

Here we assume each common neighbor has a unit of resource and will equally distribute among all its neighbors.

**(5) Katz Index**. The Katz index is based on a weighted sum over the collection of all paths connecting nodes $i$ and $j$:

(S5) $$s_{ij} = \sum_{l=1}^{\infty} \beta^l (A^l)_{ij},$$

where $\beta$ is a damping factor that gives the shorter paths more weights, and $A$ is the adjacency matrix of the network. The $N \times N$ similarity matrix $S = (s_{ij})$ can be written in a compact form [68]:

(S6) $$S = (I - \beta A)^{-1} - I,$$

where $I$ is the identity matrix. The damping factor $\beta$ is a free parameter and should be lower than the reciprocal of the absolute value of the largest eigenvalue $|\lambda_{\max}|$ of $A$ (in our calculations, we choose $\beta = 0.5/|\lambda_{\max}|$).

**(6) Average Commute Time**. The average commute time index is motivated by a random walk process on the network [62]:

(S7) $$s_{ij} = \frac{1}{l_{ii}^{+} + l_{jj}^{+} - 2l_{ij}^{+}},$$

where $l_{ij}^{+}$ is the entry of pseudoinverse of the Laplacian matrix $L \equiv D - A$, where $D = \mathrm{diag}\{k_1, k_2, \cdots, k_N\}$ is the degree matrix and $A$ is the adjacency matrix.

**(7) Network embedding**. Network embedding aims to represent each node $i \in \mathcal{V}$ into a low-dimensional space $\mathbb{R}^d$ by using the proximity between nodes [69]. After embedding the nodes in the network into a low-dimensional space, we can directly calculate the distance between any two nodes in the transformed space, which can then be used in many downstream analysis, such as multi-label classification, networks reconstruction as well as link prediction. There are many existing network ambedding methods, such as Structural Deep Networks Embedding

(SDNE) [70], LINE [71], DeepWalk [72], GraRep [73], and LE [74]. Though network embedding methods can efficiently perform link prediction for large-scale networks, the embedding process itself will cause information loss, which might affect the performance of link prediction. In addition, for sparse network, embedding methods cannot provide the representations of isolate nodes since no attribute is available. We choose DeepWalk to compare with our method and the code is downloaded from github: https://github.com/phanein/deepwalk.

**(8) Non-negative matrix factorization**. Suppose the adjacency matrix $A$ of a network is also non-negative where each node is represented by the corresponding column. Then non-negative matrix factorization (NMF) is aiming to find two non-negative matrices $U_{N\times k}$ and $V_{k\times N}$ so that [75, 76]:

$$A = UV, \tag{S8}$$

where $N$ is the network size and $k < N$ is the dimension of latent space. Then, similar to the network embedding methods, we can calculate the distance (similarity) between any two nodes in the latent space and realize the task of link prediction [77].

Other similarity-based algorithms, such as SimRank [78], Random Walks [79], Random Walks with Restart [80], Negated Shortest Path [81] can also be used for link prediction.

1.2. **Maximum likelihood algorithms.** The maximum likelihood algorithms assume that real networks have some structure, i.e., hierarchical or community structure. The goal of these algorithms is to select model parameters that can maximize the likelihood of the observed structure.

**(9) Stochastic Block Model**. As one of the most general network models, the stochastic block model (SBM) assumes that nodes are partitioned into groups and the probability of two nodes are connected depends solely on the groups they belong to [82, 83]. The SBM assumes that a link with higher reliability has higher existent probability, and the reliability of a link is defined as [84]:

$$R_{ij}^{L} = \frac{1}{Z}\sum_{P\in\mathcal{P}}\left(\frac{l_{\sigma_i\sigma_j}^{O}+1}{r_{\sigma_i\sigma_j}+2}\right)\text{exp}[-\mathcal{H}(P)], \tag{S9}$$

where $\mathcal{P}$ represents the partition space of all possible partitions, $\sigma_i$ is the group that node $i$ belongs to the partition $P$, $l_{\sigma_i\sigma_j}^{O}$ is the number of links between groups $\sigma_i$ and $\sigma_j$ in the observed network, $r_{\sigma_i\sigma_j}$ is the maximum possible number of links between them, the function $\mathcal{H}(P) \equiv \sum_{\alpha\leq\beta}[\ln(r_{\alpha\beta}+1)+\ln\begin{pmatrix} r_{\alpha\beta} \\ l^O_{\alpha\beta}\end{pmatrix}]$, and $Z \equiv \sum_{P\in\mathcal{P}} \exp[-\mathcal{H}(P)]$. In practice, we can use the Metropolis algorithms to sample relevant partitions that significantly contribute to the sum over the partition space $\mathcal{P}$. This allows us to calculate the link reliability efficiently.

**(10) Hierarchical Structure Model**. Many real networks have hierarchical structure, which can be represented by a dendrogram $D$. One can assign a probability $p_r$ to each internal node $r$ of $D$. Then the connecting probability of a pair of leaves is given by $p_{r'}$, where $r'$ is the lowest common ancestor of these two leaves. Denote $E_r$ as the number of edges in the network whose endpoints have $r$ as their lowest common ancestor in the dendrogram $D$. Let $L_r$ and $R_r$ be the number of leaves in the left and right subtrees rooted at $r$, respectively. Then the likelihood of $D$ associated with a set of probabilities $\{p_r\}$ is given by [85]:

(S10) $$\mathcal{L}(D,\{p_r\}) = \prod_{r\in D} p_r^{E_r}(1-p_r)^{L_rR_r-E_r}.$$

For a specific $D$, the probabilities $\{\overline{p}_r\}$ that maximize $\mathcal{L}(D,\{p_r\})$ are edges between the two subtrees of $r$ that are present in the network:

(S11) $$\{\overline{p}_r\} = \frac{E_r}{L_rR_r}.$$

Evaluating the likelihood $\mathcal{L}(D,\{p_r\})$ at this maximum yields

(S12) $$\mathcal{L}(D) = \prod_{r\in D}\left[\bar{p}_r^{\bar{p}_r}(1-\bar{p}_r)^{1-\bar{p}_r}\right]^{L_rR_r}.$$

One can use the Markov chain Monte Carlo (MCMC) method to sample a large number of dendrorgams $D$ with probability proportional to their likelihood $\mathcal{L}(D)$. For each pair of unconnected leaves $i$ and $j$, we calculate the connection probability $p_{ij}$ for each $D$, and then calculate the average $\langle p_{ij}\rangle$ over all the sampled dendrograms. The $\langle p_{ij}\rangle$ value yields the existent probability of the link between nodes $i$ and $j$. For each nonexistent link or node pair $(i,j)$, we calculate the average connecting probability $\langle p_{ij}\rangle$ over all sampled dendrograms and the node pairs with highest $\langle p_{ij}\rangle$ are missing links.

Since the SBM-based link prediction method introduced in Ref [84] has demonstrated better performance than this hierarchical structure model, in this work we chose the former as a representative maximum likelihood algorithm to compare with our DGM-based link prediction method.

##### 1.3. **Other algorithms.**

**(11) Matrix Completion**. The goal of matrix completion is to recover a low-rank matrix $L$ from a large matrix $A$, which can be used to infer the missing links of a network. The matrix $L$ can be calculated by solving the convex optimization problem [86]:

$$\min_{L,S} \|L\|_* + \lambda|S|_1, \text{s.t. } A = L + S. \tag{S13}$$

Here, $S$ is a sparse matrix. $\|\cdot\|$ denotes the nuclear norm of a matrix, and $|\cdot|$ represents the sum of the absolute values of each matrix element and $\lambda$ is a positive parameter. The matrix $S$ is defined as follows: if two nodes $(i, j)$ are connected in the observed network, then the score $s_{ij} = A_{ij}$; otherwise, $s_{ij} = L_{ij}$. Note that this method works under a strong assumption that the number of entries in matrix $A$ should be sufficiently large to complete the low-rank matrix [87]. Therefore, it is only suitable for specific networks rather than general complex networks.

**(12) Graph Convolutional Neural Network**. Graph Convolutional Neural Network (GCN) is a variant of convolutional neural networks (CNNs) that operate directly on graphs [88]. Recently, a new link prediction method: SEAL (learning from Subgraphs, Embeddings, and Attributes for Link prediction) was developed [89]. This method combines graph structure features, latent features, and explicit features into a single GCN. In particular, the input to GCN is a local subgraph around each target link. Those local subgraphs capture graph structure feature related to link existence. The latent and explicit features can be naturally combined in GCN by concatenating node embedding and node attributes in the node information matrix for each subgraph. The SEAL code can be downloaded from github: https://github.com/muhanzhang/SEAL.

There are also many probabilistic model based link prediction methods (which typically have high time complexity [90]), such as Probabilistic Relational Model [91], Probabilistic Entity Relationship Model [92], and Stochastic Relational Model [93], as well as many other methods, such as Structural Perturbation Method [94], etc. For a more complete list of existing link prediction methods, please see the review articles [90, 95]. In this work, we compare our DGM-based link prediction method with several representative link prediction methods that have either relatively higher performance or lower time complexity.

#### 2. Performance metrics of link prediction methods

The two most commonly used metrics to quantify the performance of link prediction methods are AUC and Precision. To calculate those two metrics, we first split the total link set $\mathcal{E}$ into two parts: (i) a fraction of links as the test or probe set $\mathcal{E}_\mathrm{P}$, which will be removed from the network; and (ii) the remaining fraction of links as the training set $\mathcal{E}_\mathrm{T}$, which will be used to recover the removed links.

If we shuffle all the existing links and randomly select an $f$ fraction of links from the total link set $\mathcal{E}$ as the probe set, then those links originating from hubs (i.e., high-degree nodes) will be more likely to be selected (especially for networks with strong degree heterogeneity), while those links originating from low-degree nodes will be less likely to be selected. To avoid any bias introduced by this edge-based splitting method, we employ a node-based splitting method. See Algorithm 1 for details and Fig. S5 for a demonstration of its fairness.

**Algorithm 1** Node-based splitting method

**Input:**
The network, defined by $\mathcal{G}(\mathcal{V}, \mathcal{E})$, where $\mathcal{V}$ denotes the node set and $\mathcal{E}$ denotes the link set.

**Output:**
The probe set $\mathcal{E}_\mathrm{P}$ with a fraction of $f$ links, and training set $\mathcal{E}_\mathrm{T}$ with a fraction of $(1-f)$ links. $\mathcal{E}_\mathrm{P} \cup \mathcal{E}_\mathrm{T} = \mathcal{E}$

Set $n = 0$
**while** $n < \lceil f \times \mathcal{E} \rceil$ **do**
  Randomly select one node $i$ from $\mathcal{V}$
  **if** node $i$ has at least one link (not an isolate node) **then**
    Randomly select one of link $(i, j)$ of node $i$
    **if** $rand() < p$[1] **then**
      Select link $(i, j)$ as one of probe links, and remove it from the network
      $n = n + 1$
    **end if**
  **end if**
**end while**

2.1. **AUC.** The AUC statistic [85] is defined to be the probability that a randomly chosen link in the probe set is given a higher score by the link prediction method than that of a randomly chosen nonexistent link. If we denote $n$ as the total number of independent comparisons, and $n'$

[1]Note: $\lceil\rceil$ represents the rounds towards the nearest integer. $rand$ denotes a random number from uniform distribution. Probability $p$ does not affect the fairness of algorithm and we chose $p = 0.01$ for all of networks.

denotes the times that the score of a randomly selected link in the probe set is higher than that of the nonexistent link, then:

$$\text{AUC} = \frac{n'}{n}. \tag{S14}$$

**Remark 1.** In literature [66], a slightly different definition of AUC was used:

$$\text{AUC} = \frac{n' + 0.5n''}{n}, \tag{S15}$$

where $n''$ is the times that the score of a randomly selected link in the probe set is equal to that of a randomly chosen nonexistent link. We emphasize that with this definition of AUC, our DGM-based link prediction method still displays very robust and superior performance for both undirected and directed real-world networks (see Fig. S6). However, for some similarity-based link prediction methods, this definition of AUC sometimes leads to misleading results. For instance, in Fig. S6, we find that for some similarity-based link prediction methods, the AUC with larger training set (or equivalently, smaller probe set, say $f = 0.1$) is even lower than that with smaller training set (or larger probe set, say $f = 0.9$). This spurious performance enhancement with larger probe set is due to the term $n''$ in Eq. (S15). Consider a specific similarity-based link prediction method such as the preferential attachment (PA) method, where the similarity score of a node pair is defined as the degree product of the two nodes. As shown in Fig. S7(b), (d), with a larger probe set, the degrees of nodes in the remaining network display very few possible values (non-negative integers), and most of the nodes have degree zero. Consequently, it is very likely that the score of a randomly selected link in the probe set will be equal to that of a randomly chosen nonexistent link. In other words, as $f \to 1$, $n' \to 0$ and $n''/n \to 1$, rendering $\text{AUC} = (n' + 0.5n'')/n \to 0.5$. Note that the AUC definition in Eq. (S14) does not include the contribution of $n''$, hence in the case $f \to 1$, we will have $n' \to 0$, and naturally $\text{AUC} = n'/n \to 0$.

In addition, one can show that the calculation of AUC using Eq. (S15) is actually equivalent to the AUC calculated directly from the whole receiver operating characteristic (ROC) curve.

For example, as shown in Fig. S4, we plotted the whole ROC curves for 12 link prediction algorithms for the 12 real-world networks analyzed in the main text (with test set fraction $f = 0.1$). The AUC values are shown in Fig. S4, which are consistent with what we show in Fig. S6. Since we want to systematically demonstrate the performance of various link prediction methods with different fractions of training set, showing the whole ROC curves will make the presentation very complicated. Hence in the main text, we did not use the ROC curves to present our results.

2.2. **Precision.** To calculate the Precision of a link prediction method, we first sort the non-observed links in descending order according to their scores assigned by the link prediction method. We take the top-$L$ non-observed links as the predicted ones, and count how many of the $L$ links belong to the probe set, denoted as $L_{\rm p}$. Then the Precision is defined as [96]:

$$\text{Precision} = \frac{L_{\rm p}}{L}. \tag{S16}$$

In the main text, we choose AUC rather than Precision as the metric to quantify the performance of various link prediction methods. The reason is that the Precision metric is quite sensitive to $L$ and the total link set $\mathcal{E}$. For example, if we compare the Precisions of $L$ being $10\%$ and $20\%$ of total links as predicted ones respectively, one can find that the Precision corresponding to $L = \lceil 0.1\mathcal{E} \rceil$ is larger than that of $L = \lceil 0.2\mathcal{E} \rceil$. This would be rather counterintuitive, since with $L = \lceil 0.2\mathcal{E} \rceil$, we have chosen more links as the probe set, and the training set is smaller, which means that the link prediction performance should degrade. Therefore, the spurious high Precision is generally caused by the increase of $L$, rather than the enhancement of link prediction method [86]. If we just calculate the Precision according to Eq. (S16), with an $f = 0.3$ fraction of probe links, and we take the top-$L$ ($L = \lceil 0.3\mathcal{E} \rceil$) non-observed links as predicted ones. Note that if we use Precision as the performance metric, our method still outperforms other methods in 8 of the 12 real networks studied in this work. None of the other methods can achieve this level of robust performance across different real networks (see Table. S1).

TABLE S1: The Precision of link prediction methods on real-world networks

| Networks | DGM | CN | PA | RA | JC | KATZ | ACT | SBM | LRMC | DW | NMF | SEAL |
|---|---|---|---|---|---|---|---|---|---|---|---|---|
| Zachary karate club | **0.202 (0.078)** | 0.048 (0.043) | 0.061 (0.055) | 0.044 (0.036) | 0 (0) | 0.083 (0.043) | 0 (0) | 0 (0) | 0 (0) | 0 (0) | 0 (0) | 0 (0) |
| Terrorist association | 0.1 (0.046) | 0.063 (0.043) | 0.067 (0.029) | 0.096 (0.040) | 0 (0) | **0.137 (0.031)** | 0 (0) | 0 (0) | 0.005 (0.003) | 0 (0) | 0 (0) | 0 (0) |
| Medieval river trade | **0.035 (0.033)** | 0 (0) | 0 (0) | 0 (0) | 0 (0) | 0 (0) | 0 (0) | 0 (0) | 0 (0) | 0 (0) | 0 (0) | 0 (0) |
| Contiguous USA | **0.08 (0.044)** | 0 (0) | 0.016 (0.021) | 0.006 (0.013) | 0 (0) | 0 (0) | 0 (0) | 0 (0) | 0 (0) | 0 (0) | 0 (0) | 0.05 (0.05) |
| Internet topology (PoP) | 0.037 (0.03) | 0.026 (0.028) | 0.042 (0.042) | 0.037 (0.025) | 0 (0) | **0.053 (0.035)** | 0 (0) | 0 (0) | 0 (0) | 0 (0) | 0 (0) | 0 (0) |
| Protein-protein interactions | 0.039 (0.017) | 0.028 (0.015) | 0.02 (0.01) | 0.027 (0.016) | 0 (0) | **0.047 (0.021)** | 0 (0) | 0 (0) | 0.003 (0.002) | 0 (0) | 0 (0) | 0 (0) |
| Cat cortex | 0.16 (0.032) | | **0.24 (0.029)** | | | | | | 0 (0) | | | |
| Consulting | **0.347 (0.014)** | | 0.344 (0.028) | | | | | | 0.0023 (0.001) | | | |
| St. Martin | **0.165 (0.043)** | | 0.082 (0.019) | | | | | | 0 (0) | | | |
| Seagrass | **0.071 (0.021)** | | 0.047 (0.021) | | | | | | 0 (0) | | | |
| Sioux Falls traffic | **0.071 (0.06)** | | 0.017 (0.03) | | | | | | 0 (0) | | | |
| Cattle | **0.199 (0.045)** | | 0.086 (0.033) | | | | | | 0 (0) | | | |

For each real-world network, we first randomly remove $L$ links ($f = 0.3$, $L = \lceil 0.3\mathcal{E} \rceil$) as the predicted ones, and the remaining links form the training set, which will be used to recover the removed links. We show the mean Precision over 10 realizations and stand deviation of Precision for each method.

#### 3. Deep Generative Models

As a class of models for unsupervised learning, generative models take a training set that consists of samples drawn from some unknown distribution $p_{\text{data}}(\mathbf{x})$, learn a distribution $p_{\text{model}}(\mathbf{x})$ that we can sample from, such that $p_{\text{model}}(\mathbf{x})$ is as similar as possible to $p_{\text{data}}(\mathbf{x})$. Generative models learn $p_{\text{model}}(\mathbf{x})$ either explicitly or implicitly [97]. In the former case, we explicitly define and solve for $p_{\text{model}}(\mathbf{x})$. In the latter case, we learn a model $p_{\text{model}}(x)$ that we can sample from it but without explicitly defining it. Deep generative models (DGMs) define distributions over a set of variables organized in multiple layers [98]. In this section, we describe two of the most commonly used DGMs: Variational Autoencoders (VAE) and Generative Adversarial Networks (GANs).

3.1. **Variational Autoencoder.** Consider a dataset consisting of $N$ samples of some continuous or discrete variable $\mathbf{x}$. VAE contains a specific probability model of data $\mathbf{x}$ and latent variable $\mathbf{z}$. We can write the joint probability distribution as $p_{\boldsymbol{\theta}}(\mathbf{x}, \mathbf{z}) = p_{\boldsymbol{\theta}}(\mathbf{x}|\mathbf{z})p_{\boldsymbol{\theta}}(\mathbf{z})$, where $\boldsymbol{\theta}$ represents neural network model parameters. VAE assumes that the data are generated as follows: (i) draw a latent variable $\mathbf{z}^{(i)}$ from some prior distribution $p_{\boldsymbol{\theta}}(\mathbf{z})$; (ii) then draw the datapoint $\mathbf{x}^{(i)}$ from the likelihood $p_{\boldsymbol{\theta}}(\mathbf{x}|\mathbf{z})$.

Note that the integral of the marginal likelihood $p_{\boldsymbol{\theta}}(\mathbf{x}) = \int p_{\boldsymbol{\theta}}(\mathbf{x}|\mathbf{z})p_{\boldsymbol{\theta}}(\mathbf{z})\mathrm{d}\mathbf{z}$ is generally intractable, since one has to evaluate over all configurations of the latent variable $\mathbf{z}$. Consequently, the posterior density $p_{\boldsymbol{\theta}}(\mathbf{z}|\mathbf{x}) = \frac{p_{\boldsymbol{\theta}}(\mathbf{x}|\mathbf{z})p_{\boldsymbol{\theta}}(\mathbf{z})}{p_{\boldsymbol{\theta}}(\mathbf{x})}$ is also intractable.

VAE approximates the intractable true posterior density $p_{\boldsymbol{\theta}}(\mathbf{z}|\mathbf{x})$ by a recognition model $q_{\boldsymbol{\phi}}(\mathbf{z}|\mathbf{x})$. Since the latent variable $\mathbf{z}$ is unobserved, it can be interpreted as a *code*. Similarly, the recognition model $q_{\boldsymbol{\phi}}(\mathbf{z}|\mathbf{x})$ is called an *encoder*. For a given datapoint $\mathbf{x}$, the encoder $q_{\boldsymbol{\phi}}(\mathbf{z}|\mathbf{x})$ will produce a distribution over the possible values of the code $\mathbf{z}$ from which the datapoint $\mathbf{x}$ could have been generated. And the likelihood $p_{\boldsymbol{\theta}}(\mathbf{x}|\mathbf{z})$ will be referred to as *decoder*, because for a given code $\mathbf{z}$, it will produce a distribution over the possible corresponding values of $\mathbf{x}$.

To quantify the difference between the encoder $q_{\boldsymbol{\phi}}(\mathbf{z}|\mathbf{x})$ and the true posterior density $p_{\boldsymbol{\theta}}(\mathbf{z}|\mathbf{x})$, we use the Kullback-Leibler divergence that measures the information loss when using the encoder to approximate the true posterior (in units of nats):

$$\begin{aligned} D_{\mathrm{KL}}\left(q_{\boldsymbol{\phi}}(\mathbf{z}|\mathbf{x})||p_{\boldsymbol{\theta}}(\mathbf{z}|\mathbf{x})\right) &\equiv \int q_{\boldsymbol{\phi}}(\mathbf{z}|\mathbf{x}) \log \frac{q_{\boldsymbol{\phi}}(\mathbf{z}|\mathbf{x})}{p_{\boldsymbol{\theta}}(\mathbf{z}|\mathbf{x})} \mathrm{d}\mathbf{z} \\ &= \mathbb{E}_{\mathbf{z}\sim q_{\boldsymbol{\phi}}(\mathbf{z}|\mathbf{x})}\left[\log q_{\boldsymbol{\phi}}(\mathbf{z}|\mathbf{x}) - \log p_{\boldsymbol{\theta}}(\mathbf{x},\mathbf{z})\right] + \log p_{\boldsymbol{\theta}}(\mathbf{x}). \end{aligned} \tag{S17}$$

Hence we have

$$\begin{aligned} \log p_{\boldsymbol{\theta}}(\mathbf{x}) &= D_{\mathrm{KL}}\left(q_{\boldsymbol{\phi}}(\mathbf{z}|\mathbf{x})||p_{\boldsymbol{\theta}}(\mathbf{z}|\mathbf{x})\right) - \mathbb{E}_{\mathbf{z}\sim q_{\boldsymbol{\phi}}(\mathbf{z}|\mathbf{x})}\left[\log q_{\boldsymbol{\phi}}(\mathbf{z}|\mathbf{x}) - \log p_{\boldsymbol{\theta}}(\mathbf{x},\mathbf{z})\right] \\ &= D_{\mathrm{KL}}\left(q_{\boldsymbol{\phi}}(\mathbf{z}|\mathbf{x})||p_{\boldsymbol{\theta}}(\mathbf{z}|\mathbf{x})\right) + \mathcal{L}(\boldsymbol{\theta},\boldsymbol{\phi};\mathbf{x}) \\ &\geq \mathcal{L}(\boldsymbol{\theta},\boldsymbol{\phi};\mathbf{x}). \end{aligned} \tag{S18}$$

In the last step, we have used the fact that the KL-divergence is non-negative. The above inequality (S18) suggests that: (i) $\mathcal{L}(\boldsymbol{\theta},\boldsymbol{\phi};\mathbf{x})$ is the (variational) lower bound of the log marginal likelihood; (ii) we can minimize $D_{\mathrm{KL}}\left(q_{\boldsymbol{\phi}}(\mathbf{z}|\mathbf{x})||p_{\boldsymbol{\theta}}(\mathbf{z}|\mathbf{x})\right)$ by maximizing $\mathcal{L}(\boldsymbol{\theta},\boldsymbol{\phi};\mathbf{x})$. Note that (ii) is the key idea of VAE, since $D_{\mathrm{KL}}\left(q_{\boldsymbol{\phi}}(\mathbf{z}|\mathbf{x})||p_{\boldsymbol{\theta}}(\mathbf{z}|\mathbf{x})\right)$ is simply intractable (due to the intractable posterior density $p_{\boldsymbol{\theta}}(\mathbf{z}|\mathbf{x})$), while $\mathcal{L}(\boldsymbol{\theta},\boldsymbol{\phi};\mathbf{x})$ is tractable.

Here

$$\begin{aligned} \mathcal{L}(\boldsymbol{\theta},\boldsymbol{\phi};\mathbf{x}) &\equiv \mathbb{E}_{\mathbf{z}\sim q_{\boldsymbol{\phi}}(\mathbf{z}|\mathbf{x})}\left[\log p_{\boldsymbol{\theta}}(\mathbf{x},\mathbf{z}) - \log q_{\boldsymbol{\phi}}(\mathbf{z}|\mathbf{x})\right] \\ &= \mathbb{E}_{\mathbf{z}\sim q_{\boldsymbol{\phi}}(\mathbf{z}|\mathbf{x})}\left[\log p_{\boldsymbol{\theta}}(\mathbf{x}|\mathbf{z})\right] - D_{\mathrm{KL}}\left(q_{\boldsymbol{\phi}}(\mathbf{z}|\mathbf{x})||p_{\boldsymbol{\theta}}(\mathbf{z})\right). \end{aligned} \tag{S19}$$

Note that the first term in Eq. (S19) is the expected log-likelihood of the datapoint $\mathbf{x}$. The expectation is taken with respect to the encoder's distribution $q_{\boldsymbol{\phi}}(\mathbf{z}|\mathbf{x})$ over the latent variable $\mathbf{z}$. This term encourages the decoder $p_{\boldsymbol{\theta}}(\mathbf{x}|\mathbf{z})$ to learn to reconstruct the data. The second term is the Kullback-Leibler divergence between the encoder's distribution $q_{\boldsymbol{\phi}}(\mathbf{z}|\mathbf{x})$ and the prior distribution $p_{\boldsymbol{\theta}}(\mathbf{z})$. This term encourages small error when using $q_{\boldsymbol{\phi}}(\mathbf{z}|\mathbf{x})$ to approximate $p_{\boldsymbol{\theta}}(\mathbf{z})$.

3.2. **Generative Adversarial Networks.** GANs are based on a game theory setup, where two players (the generator and the discriminator) compete with each other [99]. Each player can be represented by a function that is differentiable with respect to both its inputs and its parameters. The generator directly produces fake samples $\mathbf{x} = G(\mathbf{z}; \boldsymbol{\theta}^{(g)})$ with $\mathbf{z}$ the input noise variable and $\boldsymbol{\theta}^{(g)}$ the parameters. Its adversary, the discriminator, attempts to distinguish between real samples drawn from the training data and fake samples generated by the generator. The discriminator takes $\mathbf{x}$ as input and uses $\boldsymbol{\theta}^{(d)}$ as parameters. It outputs a single scalar $D(\mathbf{x}; \boldsymbol{\theta}^{(d)})$, indicating the probability of $\mathbf{x}$ belonging to the real samples.

Within the GANs framework, the discriminator is trying to minimize the cost function

(S20)
$$J^{(d)}(\boldsymbol{\theta}^{(d)}, \boldsymbol{\theta}^{(g)}) = -\frac{1}{2}\mathbb{E}_{\mathbf{x}\sim\mathbb{P}_{\text{data}}(\mathbf{x})} \log D(\mathbf{x}, \boldsymbol{\theta}^{(d)}) - \frac{1}{2}\mathbb{E}_{\mathbf{z}\sim\mathbb{P}_{\mathbf{z}}(\mathbf{z})} \log \left[1 - D(G(\mathbf{z}, \boldsymbol{\theta}^{(g)}); \boldsymbol{\theta}^{(d)})\right],$$

which is the cross-entropy cost when training a standard binary classifier with a sigmoid input [100].

For the generator, we can specify its cost function in many different ways. The simpliest way is a zero-sum game, i.e.,

$$J^{(g)}(\boldsymbol{\theta}^{(d)}, \boldsymbol{\theta}^{(g)}) = -J^{(d)}(\boldsymbol{\theta}^{(d)}, \boldsymbol{\theta}^{(g)}). \tag{S21}$$

Since $J^{(g)}$ is directly tied to $J^{(d)}$, we can summarize the entire game with a value function

$$V(\boldsymbol{\theta}^{(d)}, \boldsymbol{\theta}^{(g)}) = -J^{(d)}(\boldsymbol{\theta}^{(d)}, \boldsymbol{\theta}^{(g)}). \tag{S22}$$

During the learning, each player attempts to maximize its own payoff, and at convergence

$$\boldsymbol{\theta}^{(g)*} = \arg\min_{\boldsymbol{\theta}^{(g)}} \max_{\boldsymbol{\theta}^{(d)}} V(\boldsymbol{\theta}^{(d)}, \boldsymbol{\theta}^{(g)}). \tag{S23}$$

Unfortunately, the cost function used for the generator $G$ in the above zero-sum game is only useful for theoretical analysis. In practice, it does not perform very well, because it may not provide sufficient gradient for $G$ to learn well. Indeed, in the beginning of the learning process, $G$ performs poorly and generates fake samples that can be easily rejected by $D$ with very high confidence. In this case, $\log\left[1 - D(G(\mathbf{z}, \boldsymbol{\theta}^{(g)}); \boldsymbol{\theta}^{(d)})\right]$ saturates, rendering the generator's gradient vanish. Instead of training $G$ to minimize $\log\left[1 - D(G(\mathbf{z}, \boldsymbol{\theta}^{(g)}); \boldsymbol{\theta}^{(d)})\right]$, we can train $G$ to maximize $\log D(G(\mathbf{z}, \boldsymbol{\theta}^{(g)}); \boldsymbol{\theta}^{(d)})$. In other words, $G$ aims to maximize the log-probability that $D$ makes a mistake, rather than aiming to minimize the log-probability that $D$ makes the correct prediction. The cost of the generator is then re-defined as

$$J^{(g)}(\boldsymbol{\theta}^{(d)}, \boldsymbol{\theta}^{(g)}) = -\frac{1}{2}\mathbb{E}_{\mathbf{z}\sim\mathbb{P}_{\mathbf{z}}(\mathbf{z})} \log D(G(\mathbf{z}, \boldsymbol{\theta}^{(g)}); \boldsymbol{\theta}^{(d)}). \tag{S24}$$

However, even this modified cost function can misbehave in case the discriminator is really good [101]. Such training difficulty of GANs might be due to the divergences between GANs' training objective and generator's parameters $\boldsymbol{\theta}^{(g)}$ updating direction [102]. To address this issue, the Earth-Mover (also called Wasserstein-1) distance $W(\mathbb{Q}, \mathbb{P})$ was proposed [102]. Informally, $W(\mathbb{Q}, \mathbb{P})$ defines the minimum cost of transporting mass in order to transform the distribution $\mathbb{Q}$ into the distribution $\mathbb{P}$ (where the cost is mass times transport distance). Under mild assumptions, $W(\mathbb{Q}, \mathbb{P})$ is continuous everywhere and differentiable almost everywhere. Unlike the original GANs cost function, the Wasserstein GANs (or WGANs) is more likely to provide gradients that are useful for updating the generator. Yet, the cost function in the WGANs relies on the discriminator being a $k$-Lipschitz continuous function [102, 103]. Following the setup in [102], the cost functions of the generator and the discriminator are:

$$J^{(g)}(\boldsymbol{\theta}^{(d)}, \boldsymbol{\theta}^{(g)}) = -\mathbb{E}_{\mathbf{z}\sim\mathbb{P}_{\mathbf{z}}(\mathbf{z})}[D(G(\mathbf{z}, \boldsymbol{\theta}^{(g)}); \boldsymbol{\theta}^{(d)})] \tag{S25a}$$

$$J^{(d)}(\boldsymbol{\theta}^{(d)}, \boldsymbol{\theta}^{(g)}) = -\mathbb{E}_{\mathbf{x}\sim\mathbb{P}_{\mathrm{data}}}[D(\mathbf{x}, \boldsymbol{\theta}^{(d)})] + \mathbb{E}_{\mathbf{z}\sim\mathbb{P}_{\mathbf{z}}(\mathbf{z})}[D(G(\mathbf{z}, \boldsymbol{\theta}^{(g)}); \boldsymbol{\theta}^{(d)})]. \tag{S25b}$$

The entire game with a value function $V(\boldsymbol{\theta}^{(g)}, \boldsymbol{\theta}^{(d)})$ is given by:

$$\min_{\boldsymbol{\theta}^{(g)}} \max_{\boldsymbol{\theta}^{(d)}} V(\boldsymbol{\theta}^{(g)}, \boldsymbol{\theta}^{(d)}) = \mathbb{E}_{\mathbf{x}\sim\mathbb{P}_{\mathrm{data}}(\mathbf{x})}[D(x, \boldsymbol{\theta}^{(d)})] - \mathbb{E}_{\mathbf{z}\sim\mathbb{P}_{\mathbf{z}}(\mathbf{z})}[D(G(z, \boldsymbol{\theta}^{(g)}); \boldsymbol{\theta}^{(d)})]. \tag{S26}$$

#### 4. DGM-BASED LINK PREDICTION

The key idea of our DGM-based link prediction method is to treat the adjacency matrix of a network as the pixel matrix of an image. By perturbing the original input image in $M$ different ways, we obtain a pool of perturbed images. Those perturbed images will be fed into a DGM to create $S$ fake images that look similar to the input ones. The existent likelihood of the link between nodes $i$ and $j$, denoted as $\alpha_{ij}$, in a fake network is simply given by $\alpha_{ij} = 1 - P_{ij}$, where $P_{ij}$ is the rescaled pixel value (ranging from $0$ to $1$) in its corresponding fake grayscale image. Finally, we take the average value $\langle \alpha_{ij} \rangle = 1 - \langle P_{ij} \rangle$ over all the $S$ fake images to get the overall existent likelihood of the node pair $(i, j)$. In the following, we describe the details of two DGMs, i.e., VAE and GANs in solving the link prediction problem.

4.1. **VAE-based link prediction.** We employed the Tensorflow implementation of Convolutional Variational Autoenconders [104]. In our calculation, we chose the following parameters: number of latent variables is 20, Mini-batch size for data subsampling is 128. We test the VAE-based link prediction for a directed network with $28$ nodes and $118$ links, whose adjacency matrix looks like the letter **E**. We randomly remove $5\%$ links from the total links to be the probe set, and the remaining links form the training set. Then, we perturb the network by randomly removing $5$ links in $M$ different ways to acquire $M(M = 1000)$ networks as the training input to VAE. The AUC of VAE-based link prediction is shown in Fig. S16(B), which is much lower than that of GANs-based link prediction in Fig. S16(C). Moreover, the fake images generated by VAE (Fig. S16(D)) are generally more blurry than those generated by GANs (Fig. S16(E)). This partially explains why the performance of VAE-based link prediction is worse than that of the GANs-based link prediction [100]. (See Sec. 4.2.2 for more detailed explanations.)

4.2. **GANs-based link prediction.** We implemented the Wasserstein GAN using Python 2.7 with deep learning open-source package TensorFlow [105]. The stride size for both generator and discriminator is $2 \times 2$. The number of convolutional and deconvolutional kernels are $128$ and $64$ for generator and discriminator, respectively. Being affected by convolutional (deconvolutional) layer as well as the stride of filters, the network size should be divisible by $4$, hence we add some isolate nodes (at most 3) to each network, while those nodes are excluded in the calculating of AUC. We use Mini-batch gradient descent with mini-batch size of $32$ to perform optimization. The learning rate (used to control the speed in adjusting the weights of neural network of loss gradient) is set to be $\eta = 0.0002$, clipping parameter (used to clip the value of weights) $c = 10^{-7}$. In order to achieve stability, the discriminator is trained two steps and then alternatively training the generator for one step (see Algorithm 2 for training process of WGAN in link prediction).

4.2.1. *Selection of hyperparameters.* Note that the AUC of GANs-based link prediction can be affected by the following hyperparameters: (1) total epochs $T$ (one epoch in stochastic gradient descent is defined as a single pass of training data through neural networks); (2) size of the input pool $M$; (3) perturbation strategy, including randomly adding or removing links; (4) fraction of perturbed links (adding $a$ links or removing $r$ links); and (5) final epoch window $(T - \Delta T, T)$ selected to calculate the average AUC.

For all of the networks (if not specified), we chose $T = 150$, $M = 1000$, $\Delta T = 20$. We systematically examined these hyperparameters in the link prediction of a directed network whose adjacency matrix looks similar to the letter **E**. We found that: (i) the larger input pool can render the AUC converge to maximum value faster, while the average AUC over final $\Delta T$ epochs is lower (Fig. S8(a), (d)); (ii) randomly removing a few links facilities the AUC of links prediction (Fig. S8(b), (e)), while randomly adding links is always inducing lower AUC (Fig. S8(c), (f)). Interestingly, for $r = 0$ (i.e., without introducing any perturbation to the network), the AUC is still very high. But we do see larger fluctuations than that of $r = 5$ in the final epoch window. Fig. S8(g)-(l) shows similar results for an undirected network (random graph).

In total, the results presented in Fig. 4 and Fig. 5 of main text are conservative estimation of the performance of our GANs-based link prediction, because we didn't fine tune hyperparameters for each network to achieve the best performance.

**Remark 2.** We emphasize that the framework of our GANs-based method in link prediction is significantly different from that in computer vision, where GANs are trained to produce fake images by learning input images' distributions. Such learning scheme may lead to a potential issue of GANs unable to learn the distribution of real data indeed [106]. Here, we infer the missing pixels (links) from the generated fake images (networks) with a single input image and its various random perturbations.

---

**Algorithm 2** WGANs for Link Prediction

---

**Input:** Learning rate $\eta$, clipping parameter $c$, batch size $m$, Number of iterations for discriminator per generator iteration $n_{discri}$

**Input:** Initial weights $\boldsymbol{\theta}^{(d)}$ for discriminator and $\boldsymbol{\theta}^{(g)}$ for generator

  **while** $\boldsymbol{\theta}^{(d)}$ has not converged **do**

    **for** $t = 0, ..., n_{discri}$ **do**

      *# Update parameter for Discriminator*

      Sample batch from historical data:

      $\{x^{(i)}\}_{i=1}^{m}\ from\ \mathbb{P}_{data}$

      Sample batch from Gaussian distribution:

      $\{z^{(i)}\}_{i=1}^{m}\ from\ \mathbb{P}_{Z}$

      Update discriminator nets using gradient descent:

      $g_{\boldsymbol{\theta}^{(d)}} \leftarrow \nabla_{\boldsymbol{\theta}^{(d)}}[-\frac{1}{m}\sum_{i=1}^{m} D(x^{(i)}) + \frac{1}{m}\sum_{i=1}^{m} D(G(z^{(i)}))]$

      $\boldsymbol{\theta}^{(d)} \leftarrow \boldsymbol{\theta}^{(d)} - \eta \cdot RMSProp^{2}(\boldsymbol{\theta}^{(d)}, g_{\boldsymbol{\theta}^{(g)}})$

      $\boldsymbol{\theta}^{(d)} \leftarrow clip^{2}(\theta^{(d)}, c, 1-c)$

    **end for**

    *# Update parameter for Generator*

    Update generator nets using gradient descent:

    $g_{\boldsymbol{\theta}^{(g)}} \leftarrow \nabla_{\boldsymbol{\theta}^{(g)}}\frac{1}{m}\sum_{i=1}^{m} D(G(z^{(i)}))$

    $\boldsymbol{\theta}^{(g)} \leftarrow \boldsymbol{\theta}^{(g)} - \eta \cdot RMSProp(\boldsymbol{\theta}^{(g)}, g_{\boldsymbol{\theta}^{(g)}})$

  **end while**

---

4.2.2. *A heuristic explanation from deep learning perspective.* Here, we briefly explain why our GANs-based link prediction works.

Denote the $N \times N$ adjacency matrix of the input network to GANs as $Y$. Here, $Y_{ij} = 1$ if there is a link from node $j$ to node $i$; otherwise $Y_{ij} = 0$. Denote $G(z)$ as the fake network generated by the generator $G$, where each element $G(z)_{ij}$ represents the existent probability of corresponding node pair $(i, j)$ in the fake network. Then, the weighted adjacency matrix of the reconstructed network, denoted as $W$, is given by:

$$W \approx G(z). \tag{S27}$$

[2]Note: $RMSProp$ denotes Root Mean Square Propagation [107] and $clip$ is the operation to clip tensor values to a specified $min$ and $max$.

Denote the ground truth of the targeted network as $R$, then the probe set and non-existent link set are given by:

$$\begin{aligned} \mathcal{E}_{\mathrm{p}} &: R_{ij} = 1, Y_{ij} = 0, \\ \mathcal{E}_{\mathrm{ne}} &: R_{ij} = 0, Y_{ij} = 0. \end{aligned} \tag{S28}$$

The AUC of our GANs-based link prediction is given by:

$$\text{AUC} = P(W_{ij} > W_{i'j'}), \text{where}(i,j) \in \mathcal{E}_{\mathrm{p}}, (i',j') \in \mathcal{E}_{\mathrm{ne}}. \tag{S29}$$

We denote $\Gamma^{ij}$ as the neighbor set of entry $(i,j)$. Then, a link $(i,j)$ belongs to one of the following cases according to the existent probability of its neighbors and itself within the input network: (i) $Y_{ij} = Y_{\Gamma^{ij}}$. For these links, the generated existent probabilities in reconstructed networks satisfy $W_{ij} \approx W_{\Gamma^{ij}} \sim 0$, if $Y_{ij} = 0$; and $W_{ij} \approx W_{\Gamma^{ij}} \sim 1$, if $Y_{ij} = 1$. (ii) $Y_{ij} = 0, Y_{\Gamma^{ij}} = 1$. Following the training process described in Section. 3.2, the generator $G$ of GANs is trained to optimize its neurons' weights $\theta^{(g)}$ such that the produced fake networks are sufficiently similar to input (real) ones. Therefore, the predicted (missing) pixels tend to have similar intensities with their surrounding pixels [108]. Equivalently, the predicted existent probabilities of missing links tend to be consistent with existent probabilities of surrounding links, which can be expressed as:

$$W_{ij} = \underset{\theta^{(g)}}{\text{argmin}} \sum_{\Gamma^{ij}} [W_{ij} - W_{\Gamma^{ij}}]^2. \tag{S30}$$

Since $W_{\Gamma^{ij}} \approx Y_{\Gamma^{ij}} \sim 1$, to minimize Eq. (S30), $W_{ij}$ will exceed $0$, which means this unobserved link in the original network has higher existent probability in the generated fake networks and hence be one of predicted links. In other words, if the existent probabilities $W_{\Gamma^{ij}}$ of link $(i,j)$'s neighbors have higher values, then the existent probability $W_{ij}$ of link $(i,j)$ also tends to be higher in the generated networks. That is, if an unconnected nodes pair is surrounded by connected nodes pairs, then it has higher existent probability and vice versa (see Fig. S9 for two examples).

**Remark 3.** We have introduced a notation $\Gamma^{ij}$ in Eq. (S30) to represent the neighboring pixels (or surrounding links) of pixel (link) $(i,j)$. The shape of the neighborhood of pixel $(i,j)$ is determined by the dimension of filters in the convolutional layer. Hence, our GANs-based link prediction method can also generate various fake networks to represent different scales of neighborhood interactions. Such adoption of filters can be applied in the link prediction by tuning the dimension of filters.

4.2.3. *A heuristic explanation from network perspective.* One of the big advantages of deep neural networks is that they can extract the features within the input data. Here, our key idea of DGM-based link prediction is to treat the adjacency matrix of a network as the pixel matrix of an image. Therefore, given a network, expressed by an adjacency matrix, or equivalently the pixel matrix, GANs can extract the features, denoting as a small $m \times n$ matrix, of the input pixel matrix by filters. Taking a feature example with $m = 2, n = 4$ at the $i$-th row and $j$-th column in the adjacency matrix:

$$\begin{pmatrix} a_{11} & a_{12} & a_{13} & \cdots & a_{1j} & a_{1,j+1} & a_{1,j+2} & a_{1,j+3} & \cdots & a_{1N} \\ a_{21} & a_{22} & a_{23} & \cdots & a_{2j} & a_{2,j+1} & a_{2,j+2} & a_{2,j+3} & \cdots & a_{2N} \\ \vdots & \vdots & \vdots & \ddots & \vdots & \vdots & \vdots & \vdots & \ddots & \vdots \\ a_{i1} & a_{i2} & a_{i3} & \cdots & 1 & 1 & 0 & 1 & \cdots & a_{iN} \\ a_{i+1,1} & a_{i+1,2} & a_{i+1,3} & \cdots & 1 & 1 & 1 & 1 & \cdots & a_{i+1,N} \\ \vdots & \vdots & \vdots & \ddots & \vdots & \vdots & \vdots & \vdots & \ddots & \vdots \\ a_{N1} & a_{N2} & a_{N3} & \cdots & a_{Nj} & a_{N,j+1} & a_{N,j+2} & a_{N,j+3} & \cdots & a_{NN} \end{pmatrix}. \tag{S31}$$

We can define the similarity between two nodes $(u, v)$ by only considering the topological information within the feature (highlighted as a red submatrix in Eq. (S31)):

$$s_{uv} = |\Gamma(u) \cap \Gamma(v)|. \tag{S32}$$

We take the node pair $(i, i + 1)$ as an example, then within the red feature submatrix we have $\Gamma(i) = \{j, j + 1, j + 3\}$ and $\Gamma(i + 1) = \{j, j + 1, j + 2, j + 3\}$, then we have $s_{i,i+1} = 3$, i.e., node $i$ and node $i + 1$ have three common neighbors (within the feature submatrix). Here, we conclude that node $i$ is similar to node $i + 1$, indicating that if there is a link from node $j + 2$ to node $i + 1$, then with high probability there exists a link from node $j + 2$ to $i$. Note that for a traditional similarity-based method, i.e., common neighbors (CN), it will conclude that with high probability there is a link from node $i$ to $i + 1$, quite different from what will be predicted from our method.

4.2.4. *Time complexity of GANs-based link prediction.* For the DGM, we leverage Generative Adversarial Networks (GANs) that consist of two deep artificial neural networks: generator and discriminator. Both are convolutional neural networks. According to the Ref [109], the total time complexity of all convolutional layers is:

$$O(\sum_{i=1}^{d} n_{l-1} \cdot {s_l}^2 \cdot n_l \cdot m_l^2), \tag{S33}$$

where $l$ is the index of a convolutional layer, $d$ is the number of convolutional layers. $n_l$ is the number of filters in the $i$-th layer and $n_{l-1}$ is the number of input channels of the $l$-th layer. $s_l$ is the spatial size of the filter and $m_l$ is the spatial size of the output feature map. For all of networks, we choose the parameters as: $d = 2$, $n_1 = 128$, $n_2 = 64$, $s_{1,2} = 5$, $m_1 \approx N/2$ and $m_1 \approx N/4$, where $N$ is the network size (the number of nodes). We only have $1$ channel, so the number of input channels for all of layers is always $1$. Then the time complexity to train the GANs of a give network is proportional to $N^2$.

#### 5. NODE RELABELING ALGORITHM

Apparent, Our DGM-based link prediction relies on the visual patterns within the network. As shown in Fig. S10 for synthetic networks generated by stochastic block model (SBM), with a fixed density of between-cluster, the performance of our DGM method gradually improves with increasing in-cluster density and stronger modular structure. By contrast, with a fixed in-cluster density, the performance of our DGM method deteriorates with increasing between-cluster density and weaker modular structure. To reach the optimal performance of our DGM-based link prediction, one should leverage any existing structural features in the network and label the nodes accordingly. After node labeling, those structural patterns naturally emerge in the matrix (image) presentation. Then, DGM is able to extract the most important structural patterns of the network as the key features of the corresponding image. Node labeling can be achieved in different ways.

5.1. **Node relabeling algorithm based on Louvain community detection.** Community detection aims to partition a network into communities of densely connected nodes, while nodes belonging to different communities are connected sparsely. For undirected networks, the quality of the partition can be measured by modularity [110, 111]:

$$Q = \frac{1}{2w}\sum_{i,j}[w_{ij} - \frac{w_i w_j}{2w}]\delta(c_i, c_j), \tag{S34}$$

where $w_{ij}$ is the weighted adjacency matrix of the network, $w_i = \sum_j w_{ij}$ is the sum of the edge weights connected to node $i$, $c_i$ is the community index to which node $i$ is assigned, $\delta(m, n) = 1$ if $m = n$ otherwise $\delta(m, n) = 0$ and $m = \frac{1}{2}\sum_{ij} w_{ij}$.

Ref [111] introduced an algorithm to find the partition with high modularity. This is often called the Louvain algorithm, which consists of two iterative phases. (i) Each node is assigned into different communities initially, then calculate the corresponding gain of modularity by removing node $i$ from its community and place it in the community of its neighbor $j$. Place $i$ in the community with maximum gain ($i$ will stay in its original community if the gain is negative). Repeat this process for all nodes until no further improvement can be achieved; (ii) Build a new network within which the nodes are the communities obtained in the first phase. The links between nodes with the same community are self-loops in the new network, while the link weight between new nodes is the sum weight of links between nodes in the corresponding two communities. Reapply the procedure in phase (i) to the weighted network obtained from phase (ii). Iterate the process until reaching the maximum modularity. In this work, we employ the Louvain algorithm implemented in Python package NetworkX 1.10 [112]. For each undirected network, this algorithm assigns each node into a community, then we relabel the nodes within the same community with similar labels.

5.2. **Node relabeling algorithm based on multiple resolution modular detection.** For directed network, we used the approach proposed in Ref [113], which can detect the modular structure for multiple resolution by introducing a resistance factor $r$. The modularity for directed network $w_{ij}$ after rescaled by $r$ is given by:

$$Q_r = \frac{1}{2w'} \sum_i \sum_j (w'_{ij} - \frac{w'^{\text{out}}_i w'^{\text{in}}_j}{2w'}) \delta(C_i, C_j), \tag{S35}$$

where $2w' = 2w + Nr$, $w'^{\text{out}}_i = w^{\text{out}}_i + r$, $w'^{\text{in}}_j = w^{\text{in}}_j + r$. $w'_{ij} = w_{ij}$ if $i \neq j$; otherwise $w'_{ij} = r$. $w = \frac{1}{2} \sum_{ij} w_{ij}$. Then, the modular structure is detected by maximizing $Q_r$ using, for example, Tabu heuristic optimization [114]. We used the Community Detection Toolbox [115] to detect the modular structure in directed networks. For each real directed network, we range scale factor $r$ and select a partition corresponding to moderate number of modules. Finally, the nodes within the same module are assigned with similar labels.

5.3. **Consensus-based node relabeling algorithm.** In the main text, we have shown the AUC of our GANs-based link prediction in both model and real-world networks by using community-based node relabeling method. Of course, community-detection based node relabeling is just one approach that can better demonstrate the pattern in the adjacency matrix of a network. Here, we develop a consensus-based relabeling method (see Fig. S11 for an example) that generally performs better than the community-based method (see Fig. S12). In the following, we describe the details of our consensus-based node relabeling algorithm.

For given labels of nodes, the adjacency matrix of a network can be treated as a pixel matrix of an image. To quantify the compactness of the image, we introduce a metric $E$ that indicates the "consensus" of pixels in each row of the pixel matrix. Formally, $E$ is defined as:

$$E \equiv \sum_{i=1}^{N} \frac{\sum_{\alpha=1}^{k_i-1} |\Gamma(i)_{\alpha+1} - \Gamma(i)_{\alpha} - 1|}{k_i - 1}, \text{ s.t. } k_i > 1, \tag{S36}$$

where $\Gamma(i)$ is the neighbor set of node $i$, $\Gamma(i)_\alpha$ is the $\alpha$-th neighbor of node $i$, $k_i = |\Gamma(i)|$ is the degree of node $i$. To achieve the global minimum $E^*$, we adopt the simulated annealing procedure [116]: (1) choose two nodes at random with uniform probability; (2) switch the labels of the two nodes and calculate $E$ for the new adjacency matrix; (3) accept the new configuration with probability

$$p = \begin{cases} 1, & \text{if } \Delta E \leq 0 \\ e^{-\beta \Delta E}, & \text{if } \Delta E > 0, \end{cases} \tag{S37}$$

where parameter $\beta$ is the inverse temperature; (4) repeat the step (1)-(3) for total $T = 50000$ times and gradually decrease $\beta$ from $1$ to $0.05$. Finally, we choose the configuration corresponding to the minimum $E$.

5.4. **Other node relabeling algorithms.** Besides node labeling methods based on community detection or consensus, we can also leverage various matrix reordering algorithms to reveal structural patterns in the adjacency matrix of a network, such as block pattern, off-diagonal block pattern, line/star pattern, bands pattern, noise anti-pattern and bandwidth anti-pattern. In fact, matrix reordering has many applications, such as in archaeology and anthropology; cartography, graphics, and information visualization; sociology and sociometry; psychology and psychometry; ecology; biology and bioinformatics; cellular manufacturing; and operations research [117, 118]. Consequently, there exist many matrix reordering algorithms, such as Robinsonian [119, 120], Spectral [121, 122], Dimension Reduction [123, 124], Heuristic Approaches [125, 126], Graph Theoretic [127, 128], Biclustering [129] , Interactive User-Controlled [130], Cuthill-McKee algorithm [131], and Barycenter based permutation [132]. We also compared the performance our DGM-based method on a real social network (Zachary karate club) labeled by the Cuthill-McKee algorithm, the Louvain method and our own consensus-based method. We found that all node labeling methods yield very similar performance (see Fig. S13).

#### 6. Brief description of real dataset

TABLE S2: Description of each real network

| Name | N | L | Type |
|---|---|---|---|
| Zachary karate club [133] | 34 | 78 | Undirected |
| Terrorist association [134] | 62 | 152 | Undirected |
| Medieval river trade [135] | 39 | 53 | Undirected |
| Contiguous USA [136] | 49 | 107 | Undirected |
| Internet topology (PoP) [137] | 41 | 63 | Undirected |
| Protein-protein interactions [138] | 104 | 236 | Undirected |
| Facebook-NIPS [139] | 2888 | 2981 | Unirected |
| Cat cortex [140] | 52 | 820 | Directed |
| Consulting [141] | 46 | 879 | Directed |
| St. Martin [142] | 45 | 224 | Directed |
| Seagrass [143] | 49 | 226 | Directed |
| Sioux Falls traffic [144] | 24 | 76 | Directed |
| Cattle [145] | 28 | 217 | Directed |
| Little rock [146] | 183 | 2494 | Directed |
| Facebook wall posts [147, 148] | 46952 | 876993 | Directed |
| Google+ [149, 139] | 23628 | 39242 | Directed |
| US airports [150] | 1574 | 28236 | Directed |

The networks of Cat cortex, Consulting network and Sioux Falls traffic, Cattle are weighted, here we treat them as an unweighted ones. The protein-protein interaction network is a sub-network of the *S. cerevisiae* protein-protein interaction network, induced by proteins annotated with the function 'Cellular Communication' of the Gene Ontology (GO) database.

##### 6.1. List of real networks.

##### 6.2. Description of each real network.

(1) **Zachary karate club.** The social network of a karate club. Each node represents an individual and each link represents that two individuals are friends outside the club activities. The network is downloaded from: https://icon.colorado.edu/#!/networks.

(2) **Terrorist association.** The terrorist communication network from the attacks on the United States on September 11, 2001. Nodes represent the terrorists and links represent the communication between them. The network is downloaded from: http://www.orgnet.com/hijackers.html.

(3) **Medieval river trade.** The network of medieval trade and communication along the rivers of Russia. Each node represents a route or place, and each link represents the trading between two routes. The network is downloaded from: https://icon.colorado.edu/#!

/networks.

(4) **Contiguous USA.** A network of contiguous states and the District of Columbia of the USA. Nodes represent states and a link means that the two states share a land-based geographic border. The network is downloaded from: http://konect.uni-koblenz.de/networks/contiguous-usa.

(5) **Internet topology (PoP level).** Internet at the Point of Presence (PoP) level. Nodes represent routers and links mean that the message can transmit along them. The network is downloaded from: http://www.topology-zoo.org/dataset.html.

(6) **Protein-protein interactions.** A sub-network of *S. cerevisiae* protein-protein interaction network, induced by proteins annotated with the function 'Cellular Communication' of the Gene Ontology (GO) database. The network is downloaded from: http://math.bu.edu/people/kolaczyk/datasets.html.

(7) **Cat cortex.** The brain network of cat cortex, where a node represents a cortical area and a link represents large cortical tracts or functional associations. The network is downloaded from: https://sites.google.com/site/bctnet/datasets.

(8) **Consulting.** A social network of consulting company. Each node represents a employee and each link represents the frequency of information or advice requests. The network is downloaded from: https://toreopsahl.com/datasets/#Cross_Parker.

(9) **St. Martin.** A network of carbon-flow among species in St. Marks National Wildlife Refuge. Each node represents a species and each link denotes a carbon flow from one species to another. The network is downloaded from: http://cosinproject.eu/extra/data/foodwebs/WEB.html.

(10) **Seagrass.** The food web that represents the predatory interactions among species in a seagrass ecosystem. Each node represents a species and each link represents a predatory interaction. The network is downloaded from: http://cosinproject.eu/extra/data/foodwebs/WEB.html.

(11) **Sioux Falls traffic.** A network of traffic flow between intersections. Nodes represent intersections and a link represents the traffic flow between two intersections. The network is downloaded from: https://github.com/bstabler/TransportationNetworks.

(12) **Cattle.** The network of dominance behaviors observed between dairy cattles at the Iberia Livestock Experiment Station in Jenerette, Louisiana. Each node represents a cattle and a link

represents dominance of the left cattle over the right cattle. The network is downloaded from: http://konect.uni-koblenz.de/networks/moreno_cattle.

(13) **Little rock.** The food web of the Little Rock Lake. Each node represents a species and each link represents a predatory interaction. The network is downloaded from: http://konect.uni-koblenz.de/networks/maayan-foodweb.

(14) **Facebook wall posts.** A network of subset of posts to other user's wall on Facebook. Each node represents a Facebook user and each link represents a wall-post interaction.
The network is downloaded from:
http://konect.uni-koblenz.de/networks/facebook-wosn-wall.

(15) **Google+.** The social network of Google+ users. Each node represents a user and a link means that one user has the other user in his or her circle. The network is downloaded from: http://konect.uni-koblenz.de/networks/ego-gplus.

(16) **Facebook-NIPS.** The user-user friendship network. Each node represents a user and a link indicates that the node is a friend of another. The network is downloaded from: http://konect.uni-koblenz.de/networks/ego-facebook.

(17) **US airports.** The network of flights between US airports in 2010. Each node represents an airport and each link represents a connection from one airport to another. The network is downloaded from: http://konect.uni-koblenz.de/networks/opsahl-usairport.

#### 7. Supplementary References

[56] D. Liben-Nowell, J. Kleinberg, *Journal of the Association for Information Science and Technology* **58**, 1019 (2007).

[57] R. N. Lichtenwalter, J. T. Lussier, N. V. Chawla, *Proceedings of the 16th ACM SIGKDD international conference on Knowledge discovery and data mining* (ACM, 2010), pp. 243–252.

[58] B. Barzel, A.-L. Barabási, *Nature biotechnology* **31**, 720 (2013).

[59] M. Al Hasan, V. Chaoji, S. Salem, M. Zaki, *SDM06: workshop on link analysis, counter-terrorism and security* (2006).

[60] R. R. Sarukkai, *Computer Networks* **33**, 377 (2000).

[61] L. Lü, C.-H. Jin, T. Zhou, *Physical Review E* **80**, 046122 (2009).

[62] W. Liu, L. Lü, *EPL (Europhysics Letters)* **89**, 58007 (2010).

[63] T. Zhou, *et al.*, *Proceedings of the National Academy of Sciences* **107**, 4511 (2010).

[64] P. Jaccard, *Bull Soc Vaudoise Sci Nat* **37**, 547 (1901).

[65] A.-L. Barabási, R. Albert, *Science* **286**, 509 (1999).

[66] T. Zhou, L. Lü, Y.-C. Zhang, *The European Physical Journal B-Condensed Matter and Complex Systems* **71**, 623 (2009).

[67] Q. Ou, Y.-D. Jin, T. Zhou, B.-H. Wang, B.-Q. Yin, *Physical Review E* **75**, 021102 (2007).

[68] L. Katz, *Psychometrika* **18**, 39 (1953).

[69] S. T. Roweis, L. K. Saul, *Science* **290**, 2323 (2000).

[70] D. Wang, P. Cui, W. Zhu, *Proceedings of the 22nd ACM SIGKDD international conference on Knowledge discovery and data mining* (ACM, 2016), pp. 1225–1234.

[71] J. Tang, *et al.*, *Proceedings of the 24th International Conference on World Wide Web* (International World Wide Web Conferences Steering Committee, 2015), pp. 1067–1077.

[72] B. Perozzi, R. Al-Rfou, S. Skiena, *Proceedings of the 20th ACM SIGKDD international conference on Knowledge discovery and data mining* (ACM, 2014), pp. 701–710.

[73] S. Cao, W. Lu, Q. Xu, *Proceedings of the 24th ACM International on Conference on Information and Knowledge Management* (ACM, 2015), pp. 891–900.

[74] M. Belkin, P. Niyogi, *Neural computation* **15**, 1373 (2003).

[75] B. Long, Z. M. Zhang, P. S. Yu, *Proceedings of the eleventh ACM SIGKDD international conference on Knowledge discovery in data mining* (ACM, 2005), pp. 635–640.

[76] L. Duan, S. Ma, C. Aggarwal, T. Ma, J. Huai, *IEEE Transactions on Knowledge and Data Engineering* **29**, 2402 (2017).

[77] B. Chen, F. Li, S. Chen, R. Hu, L. Chen, *PloS one* **12**, e0182968 (2017).

[78] G. Jeh, J. Widom, *Proceedings of the eighth ACM SIGKDD international conference on Knowledge discovery and data mining* (ACM, 2002), pp. 538–543.

[79] K. Pearson, *Nature* **72**, 294 (1905).

[80] H. Tong, C. Faloutsos, J.-Y. Pan, *in:Proceedings of the 6th International Conference on Data Mining* (2006).

[81] D. Liben-Nowell, An algorithmic approach to social networks, Ph.D. thesis, Massachusetts Institute of Technology (2005).

[82] H. C. White, S. A. Boorman, R. L. Breiger, *American journal of sociology* **81**, 730 (1976).

[83] P. W. Holland, K. B. Laskey, S. Leinhardt, *Social networks* **5**, 109 (1983).

[84] R. Guimerà, M. Sales-Pardo, *Proceedings of the National Academy of Sciences* **106**, 22073 (2009).

[85] A. Clauset, C. Moore, M. E. Newman, *Nature* **453**, 98 (2008).

[86] R. Pech, D. Hao, L. Pan, H. Cheng, T. Zhou, *EPL (Europhysics Letters)* **117**, 38002 (2017).

[87] E. J. Candès, B. Recht, *Foundations of Computational mathematics* **9**, 717 (2009).

[88] T. N. Kipf, M. Welling, *arXiv preprint arXiv:1609.02907* (2016).

[89] M. Zhang, Y. Chen, *arXiv preprint arXiv:1802.09691* (2018).

[90] V. Martínez, F. Berzal, J.-C. Cubero, *ACM Computing Surveys (CSUR)* **49**, 69 (2017).

[91] N. Friedman, L. Getoor, D. Koller, A. Pfeffer, *IJCAI* (1999), vol. 99, pp. 1300–1309.

[92] D. Heckerman, C. Meek, D. Koller, *Introduction to statistical relational learning* pp. 201–238 (2007).

[93] K. Yu, W. Chu, S. Yu, V. Tresp, Z. Xu, *Advances in neural information processing systems* (2007), pp. 1553–1560.

[94] L. Lü, L. Pan, T. Zhou, Y.-C. Zhang, H. E. Stanley, *Proceedings of the National Academy of Sciences* **112**, 2325 (2015).

[95] Y. Hulovatyy, R. W. Solava, T. Milenković, *PloS one* **9**, e90073 (2014).

[96] L. Lü, T. Zhou, *Physica A: statistical mechanics and its applications* **390**, 1150 (2011).

[97] I. Goodfellow, Y. Bengio, A. Courville, Y. Bengio, *Deep learning*, vol. 1 (MIT press Cambridge, 2016).

[98] Z. Hu, Z. Yang, R. Salakhutdinov, E. P. Xing, *arXiv preprint arXiv:1706.00550* (2017).

[99] I. Goodfellow, *et al.*, *Advances in neural information processing systems* (2014), pp. 2672–2680.

[100] I. Goodfellow, *arXiv preprint arXiv:1701.00160* (2016).

[101] M. Arjovsky, L. Bottou, *arXiv preprint arXiv:1701.04862* (2017).

[102] M. Arjovsky, S. Chintala, L. Bottou, *arXiv preprint arXiv:1701.07875* (2017).

[103] I. Gulrajani, F. Ahmed, M. Arjovsky, V. Dumoulin, A. C. Courville, *Advances in Neural Information Processing Systems* (2017), pp. 5769–5779.

[104] Convolutional variational autoencoder – Jason Ramapuram (2016).

[105] M. Abadi, *et al.*, *arXiv preprint arXiv:1603.04467* (2016).

[106] S. Arora, Y. Zhang, *arXiv preprint arXiv:1706.08224* (2017).

[107] T. Tieleman, G. Hinton, *COURSERA: Neural networks for machine learning* **4**, 26 (2012).

[108] R. Yeh, C. Chen, T. Y. Lim, M. Hasegawa-Johnson, M. N. Do, *arXiv preprint arXiv:1607.07539* (2016).

[109] K. He, J. Sun, *Proceedings of the IEEE conference on computer vision and pattern recognition* (2015), pp. 5353–5360.

[110] M. E. Newman, *Proceedings of the national academy of sciences* **103**, 8577 (2006).

[111] V. D. Blondel, J.-L. Guillaume, R. Lambiotte, E. Lefebvre, *Journal of statistical mechanics: theory and experiment* **2008**, P10008 (2008).

[112] A. Hagberg, P. Swart, D. S Chult, Exploring network structure, dynamics, and function using networkx, *Tech. rep.*, Los Alamos National Lab.(LANL), Los Alamos, NM (United States) (2008).

[113] A. Arenas, A. Fernandez, S. Gomez, *New journal of physics* **10**, 053039 (2008).

[114] F. Glover, *Computers & operations research* **13**, 533 (1986).

[115] Community detection toolbox – Athanasios (2014).

[116] S. Kirkpatrick, C. D. Gelatt, M. P. Vecchi, *Science* **220**, 671 (1983).

[117] I. Liiv, *Statistical Analysis and Data Mining: The ASA Data Science Journal* **3**, 70 (2010).

[118] M. Behrisch, B. Bach, N. Henry Riche, T. Schreck, J.-D. Fekete, *Computer Graphics Forum* (Wiley Online Library, 2016), vol. 35, pp. 693–716.

[119] W. S. Robinson, *American antiquity* **16**, 293 (1951).

[120] D. G. Kendall, *Bulletin of the International Statistical Institute* **40**, 657 (1963).

[121] H. Sternin, Statistical methods of time sequencing., *Tech. rep.*, STANFORD UNIV CALIF DEPT OF STATISTICS (1965).

[122] M. Friendly, *The American Statistician* **56**, 316 (2002).

[123] D. Harel, Y. Koren, *International symposium on graph drawing* (Springer, 2002), pp. 207–219.

[124] N. Elmqvist, T.-N. Do, H. Goodell, N. Henry, J.-D. Fekete, *Visualization Symposium, 2008. PacificVIS'08. IEEE Pacific* (IEEE, 2008), pp. 215–222.

[125] S. B. Deutsch, J. J. Martin, *Operations Research* **19**, 1350 (1971).

[126] L. Wilkinson, *The grammar of graphics* (Springer Science & Business Media, 2006).

[127] S. Sloan, *International Journal for Numerical Methods in Engineering* **23**, 239 (1986).

[128] S. Sloan, *International Journal for Numerical Methods in Engineering* **28**, 2651 (1989).

[129] J. A. Hartigan, *Journal of the american statistical association* **67**, 123 (1972).

[130] C. Perin, P. Dragicevic, J.-D. Fekete, *IEEE transactions on visualization and computer graphics* **20**, 2082 (2014).

[131] E. Cuthill, J. McKee, *Proceedings of the 1969 24th national conference* (ACM, 1969), pp. 157–172.

[132] E. Mäkinen, H. Siirtola, *Informatica (Slovenia)* **29**, 357 (2005).

[133] W. W. Zachary, *Journal of anthropological research* **33**, 452 (1977).

[134] V. E. Krebs, *Connections* **24**, 43 (2002).

[135] F. R. Pitts, *Social networks* **1**, 285 (1978).

[136] D. E. Knuth, *The Stanford GraphBase: a platform for combinatorial computing*, vol. 37 (Addison-Wesley Reading, 1993).

[137] S. Knight, H. X. Nguyen, N. Falkner, R. Bowden, M. Roughan, *IEEE Journal on Selected Areas in Communications* **29**, 1765 (2011).

[138] D. J. Hand, *International Statistical Review* **78**, 135 (2010).

[139] J. Leskovec, J. J. Mcauley, *Advances in neural information processing systems* (2012), pp. 539–547.

[140] J. Scannell, G. Burns, C. Hilgetag, M. O'Neil, M. P. Young, *Cerebral Cortex* **9**, 277 (1999).

[141] R. L. Cross, A. Parker, *The hidden power of social networks: Understanding how work really gets done in organizations* (Harvard Business Review Press, 2004).

[142] D. Baird, J. Luczkovich, R. R. Christian, *Estuarine, Coastal and Shelf Science* **47**, 329 (1998).

[143] R. R. Christian, J. J. Luczkovich, *Ecological modelling* **117**, 99 (1999).

[144] L. J. LeBlanc, E. K. Morlok, W. P. Pierskalla, *Transportation research* **9**, 309 (1975).

[145] M. W. Schein, M. H. Fohrman, *The British Journal of Animal Behaviour* **3**, 45 (1955).

[146] N. D. Martinez, *Ecological monographs* **61**, 367 (1991).

[147] Facebook wall posts network dataset – KONECT (2017).

[148] B. Viswanath, A. Mislove, M. Cha, K. P. Gummadi, *Proc. Workshop on Online Social Networks* (2009), pp. 37–42.

[149] Google+ network dataset – KONECT (2017).

[150] T. Opsahl, *online] http://toreopsahl. com/2011/08/12/why-anchorage-is-not-that-important-binary-tiesand-sample-selection (accessed September 2013)* (2011).

#### 8. SUPPLEMENTARY FIGURES

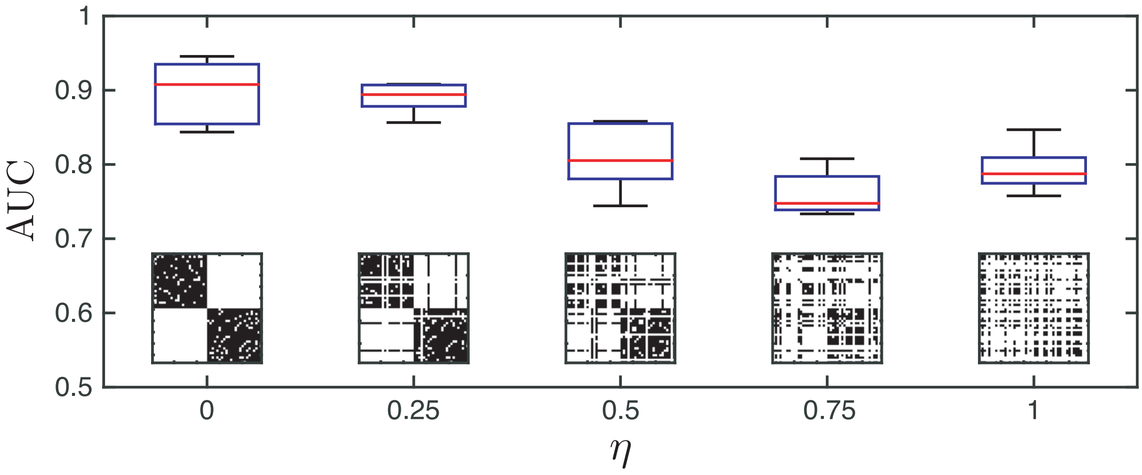

FIG. S1: **Performance of our DGM-based link prediction on a network generated by the stochastic block model.** The network has $N = 48$ nodes, and two communities (with 24 nodes for each community). The between-cluster connection probability is set to be zero and within-cluster connection probability is $0.83$. A randomly selected fraction of $f(0.25, 0.5, 0.75, 1.0)$ nodes are relabeled, rendering different ”visual patterns”. Boxplots of AUC are calculated from 10 different realizations. For each realization, we randomly divide the links into two parts: a fraction of $10\%$ links chosen as the probe set and the remaining $90\%$ links as the training set.

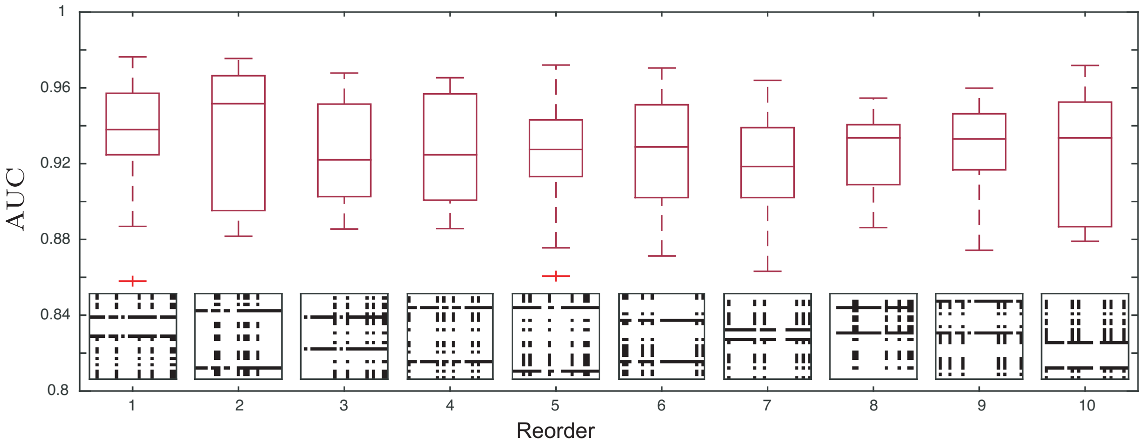

FIG. S2: **Performance of our DGM-based link prediction with different random node labelings.** We randomly label the nodes in a toy network (as shown in main text Fig.2a) in 10 different ways to obtain 10 different adjacency matrices (binary images shown in inset). Then we show the performance of our method for each adjacency matrix (labelled network) in 10 realizations using boxplots of AUC. For each realization, we randomly divide the links into two parts: a fraction of $10\%$ links chosen as the probe set and the remaining $90\%$ links as the training set. We chose $M = 1,000$ and $S = 500$ for our DGM-based method.

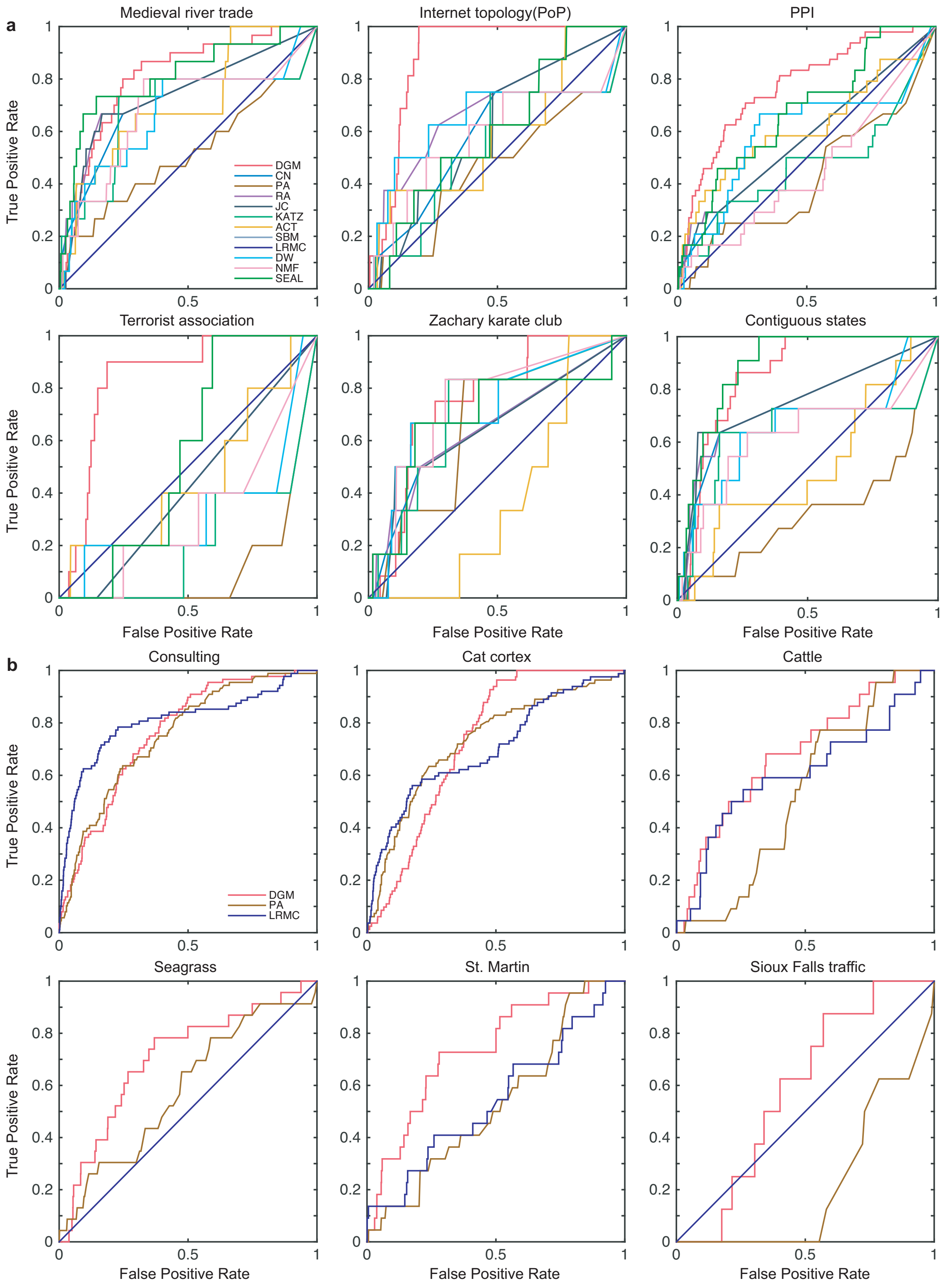

FIG. S3: **ROC curves of different link prediction methods in various real-world networks.** We randomly divide the links into two parts: a fraction of 10% links chosen as the probe set and the remaining 90% as training set.

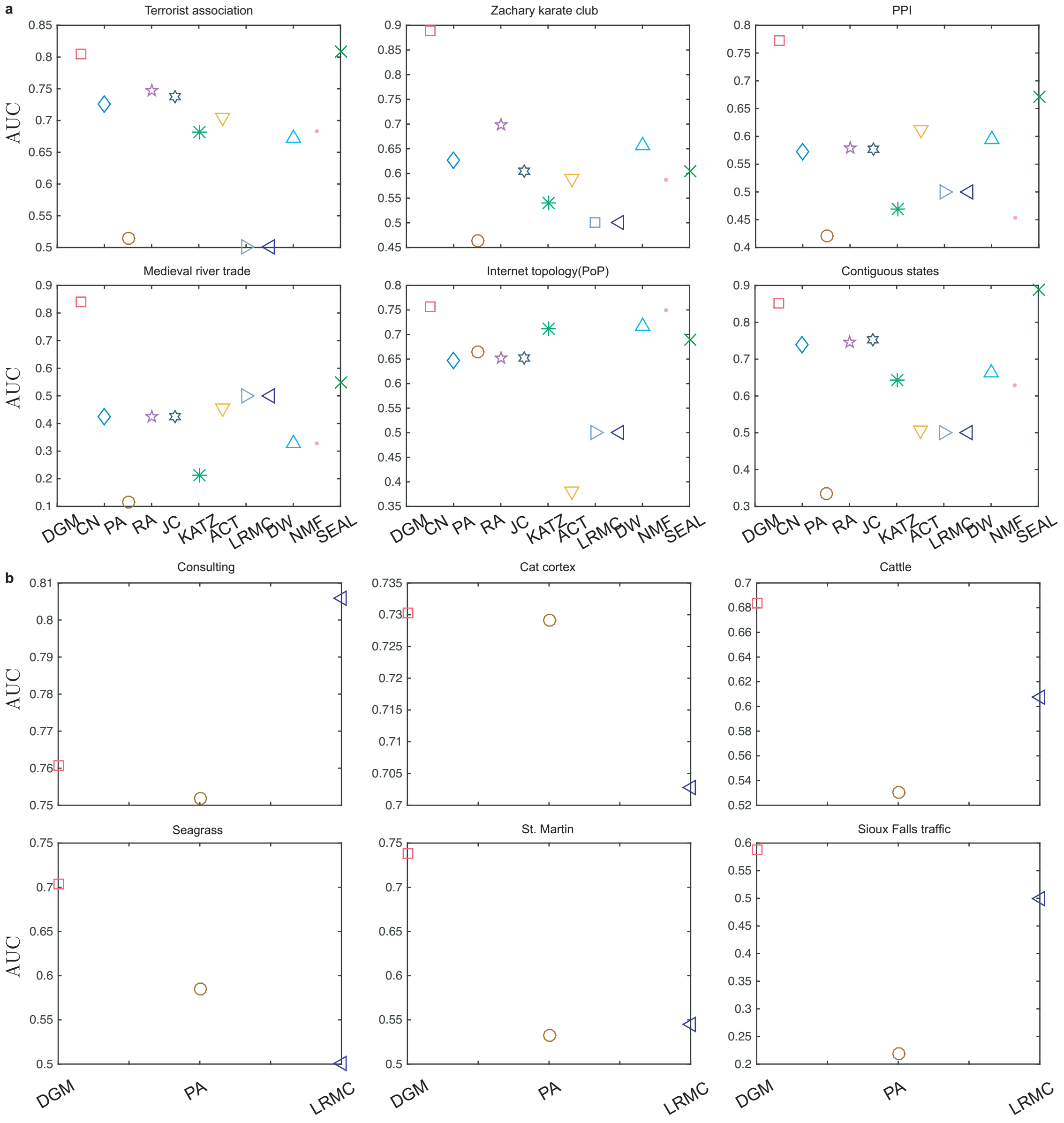

FIG. S4: **AUCs calculated directly from the ROC curves shown in Fig. S3.**

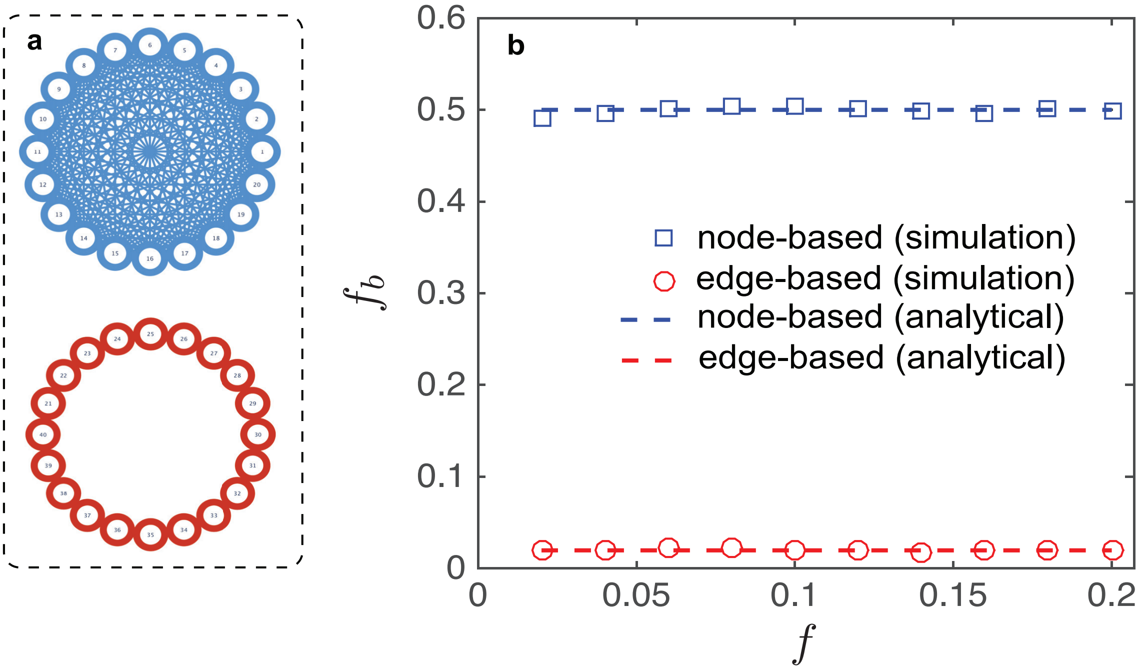

FIG. S5: **Edge-based splitting tends to select links originating from hub nodes to be part of the probe set. a,** Consider a simple network consisted of two separated subnetworks: (i) A complete graph with $N_1 = 20$ nodes; (ii) A cycle with $N_2 = 20$ nodes. The degree of each node is $k_i = N_1 - 1$ in the first subnetwork and $k'_i = 2$ in the second subnetwork. For the node-based splitting, the probability of selecting $s(s \leq N_2 - 1)$ probe links belonging to the second subnetwork is: $f_b^{\rm n} = \frac{N_2}{N_1+N_2}$. For the edge-based splitting, the probability of selecting $s$ probe links that belong to the second subnetwork is: $f_b^{\rm e} = \frac{E_2}{E_1+E_2}$, where $E_1 = \frac{N_1(N_1-1)}{2}$ and $E_2 = N_2 - 1$ are the number of total links in the first and the second subnetwork, respectively. When $N_1 = N_2 = n$, we have $f_b^{\rm n} = 0.5$, indicating the node-based splitting is perfectly fair; whereas, $f_b^{\rm e} = \frac{2}{n+2} \to 0$ for large $n$, indicating that the probability of selecting probe links originating from low-degree nodes is extremely low. **b,** The probability $f_b$ of selecting a fraction of $f$ links as probe set that belong to the second subnetwork, using the edge-based splitting (circles) or node-based splitting (squares).

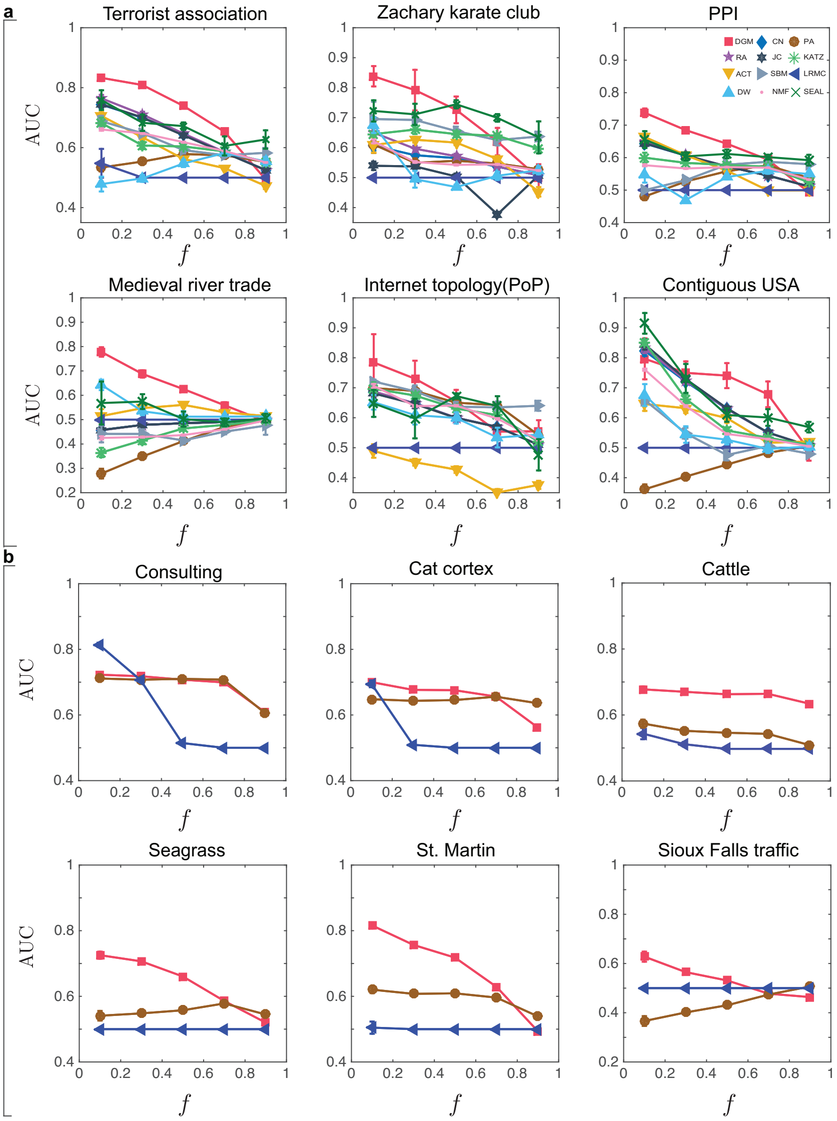

FIG. S6: **The DGM-based link prediction method displays very robust and superior performance for both undirected and directed real-world networks.** Here an alternative definition of AUC (Eq. (S15)) is used. DGM: deep generative model based link prediction; CN: common neighbors; PA: preferential attachment; RA: resource allocation; JC: Jaccard index; KATZ: Katz index; ACT: average commute time; SBM: stochastic block model; LRMC: low rank matrix completion; DW: deepwalk embedding method; NMF: non-negative matrix factorization); SEAL: learning from Subgraphs, Embeddings, and Attributes for Link prediction (see Section. 1 for details of each algorithm). **a,** Undirected networks. Top: Terrorist association network, Zachary karate club, Protein-protein interaction (PPI) network (a subnetwork of protein-protein interactions in *S. cerevisiae*). Bottom: Medieval river trade network in Russia, Internet topology (at the PoP level), Contiguous states in USA. **b,** Directed networks. Top: Consulting (a social network of a consulting company), cat cortex (the connection network of cat cortical areas), cattle (a network of dominance behaviors of cattles). Bottom: Seagrass food web, St. Martin food web, Sioux Falls traffic network. AUC of our DGM-based method is the average AUC over the last 20 epochs of the total epochs for all of networks.

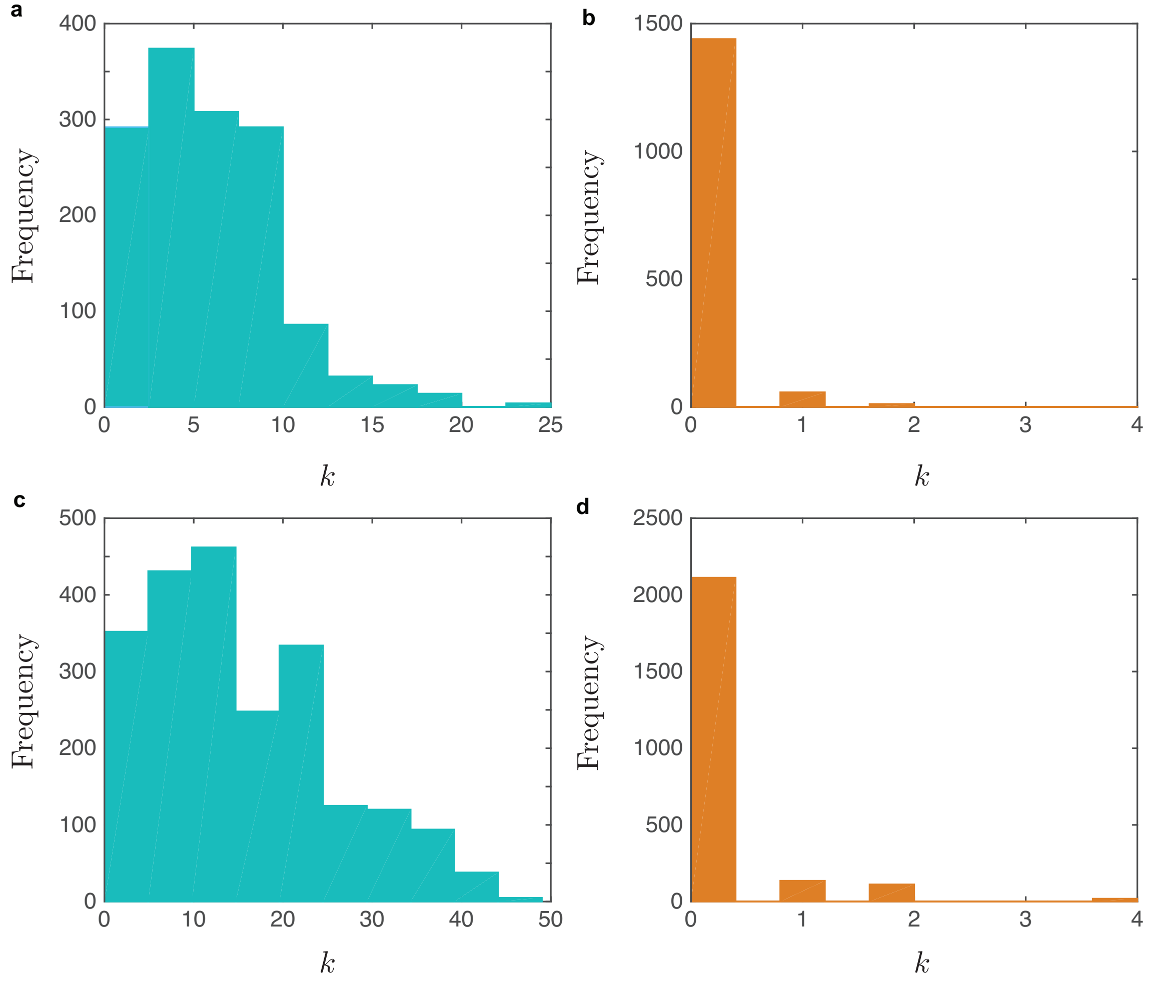

FIG. S7: **The spurious enhancement of AUC (Eq. (S15)) with larger probe set for some link prediction methods is induced by the term $n''$.** An undirected network (Medieval river trade in (**a**), (**b**)) and a directed network (Sioux Falls traffic in (**c**), (**d**)) are chosen to illustrate this point. The total links of each network are split into two parts: a fraction of $f$ as the probe set and the remaining $1-f$ as the training set. Here, we show the degree distributions of nodes in the remaining network for $f = 10\%$ (left panel) and $f = 90\%$ (right panel), respectively. For a small probe set, the possible degrees of nodes in the remaining network are very diverse ((**a**), (**c**)). Consequently, it is very unlikely that the score of a randomly selected link in the probe set will be equal to that of a randomly chosen nonexistent link. In other words, the term $n''$ in Eq. (S15) is negligible and will not affect AUC. However, for a large probe set , the degrees of nodes in the remaining network display very few possible values ((**b**), (**d**)), and most of the nodes have degree zero. Then it is very likely that the score of a randomly selected link in the probe set and that of a randomly chosen nonexistent link are both zero, rendering a large $n''$ in Eq. (S15) and hence a spuriously large AUC.

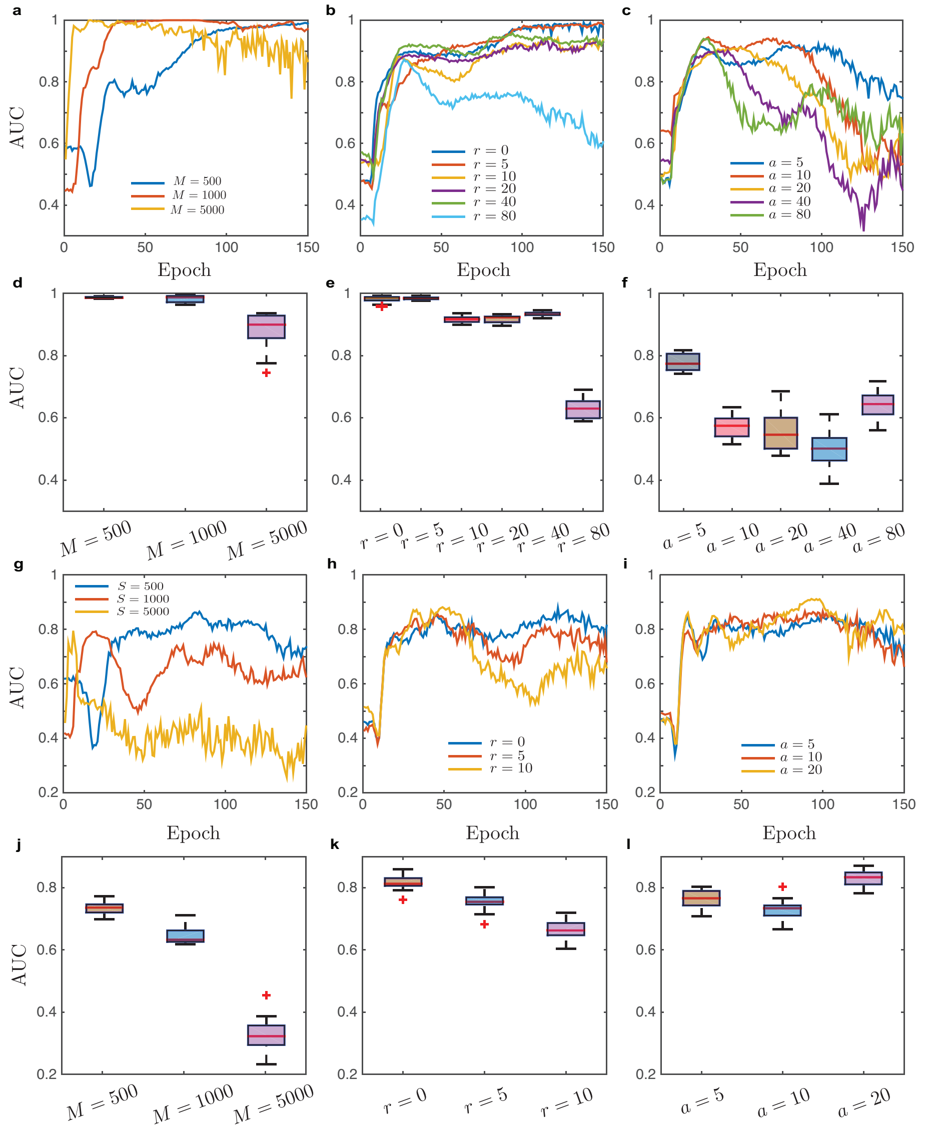

FIG. S8: **Effect of hyperparameters in GANs on the performance of link prediction.** To illustrate the effect of hyperparameters on the AUC, we perform GANs-based link prediction for on two networks. We divide the links of each network into two parts: 10% be the probe set and the remaining 90% be the training set. We perturb the training set by removing $r$ (or adding $a$) links at random in $M$ different ways to obtain the input dataset, which will be fed into the GANs. **a-f,** A directed network (of 28 nodes and 118 links) whose adjacency matrix looks like the image of letter ***E*** (with some missing pixels). AUC as a function of epoch for different input size $S$ (**a**), number of removed links $r$ (**b**), number of added links (**c**). AUC of the last 20 epochs for different input size $S$ **d**,, number of removed links $r$ (**e**), number of added links (**f**). **g-l,** An undirected random network (of 48 nodes and 40 links). AUC as a function of epoch for different input size $M$ (**g**), number of removed links $r$ (**h**), number of added links (**i**). AUC of the last 20 epochs for different input size $S$ (**j**), number of removed links $r$ (**k**), number of added links (**l**).

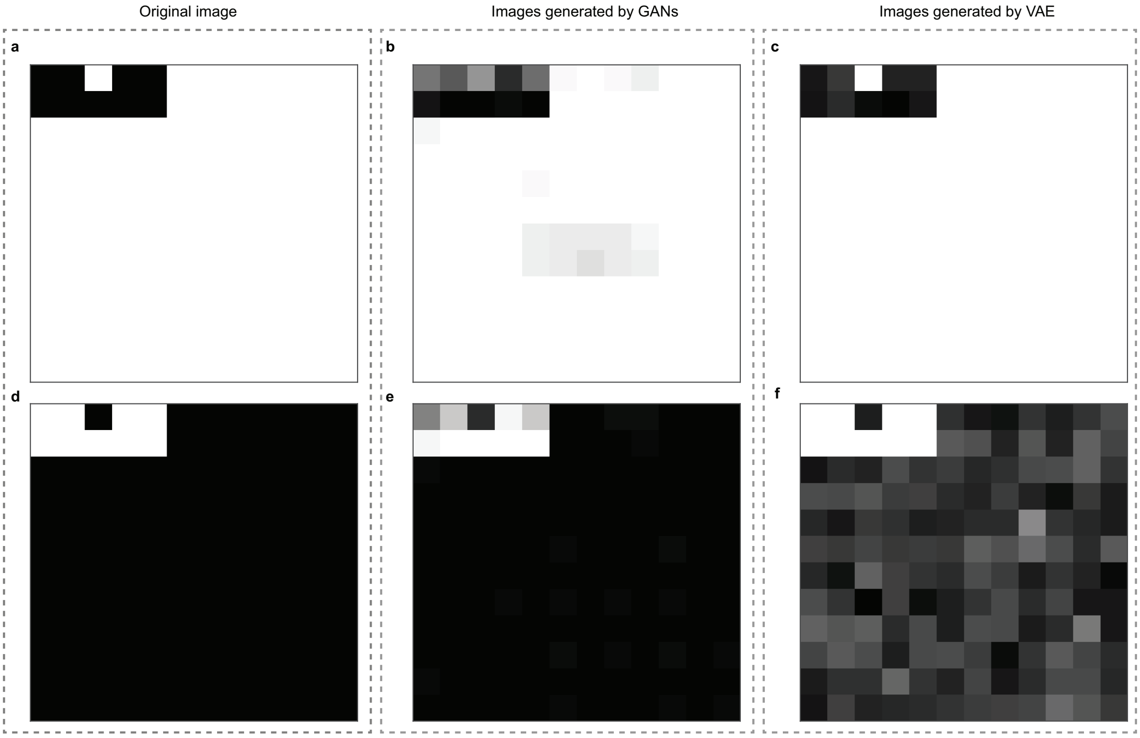

FIG. S9: **Intuitive explanation of DGM-based link prediction methods.** **a,** An missing link $(1, 3)$ is surrounded by several observed links in the original network. **b,** The snapshot of the recovered network at Epoch $= 50$ by GANs. The existent probability of this missing pixel (link) generated by GANs tends to be consistent with its surrounded pixels (links). **c,** The snapshot of the recovered network at Epoch $= 50$ by VAE. **d,** An observed link $(1, 3)$ is surrounded by several missing links in the original network. **e,** The snapshot of the recovered network at Epoch $= 50$ by GANs. The existent probabilities of several missing pixels (links) tend to be consistent with this observed pixel (link). **f,** The recovered images at Epoch $= 50$ by VAE.

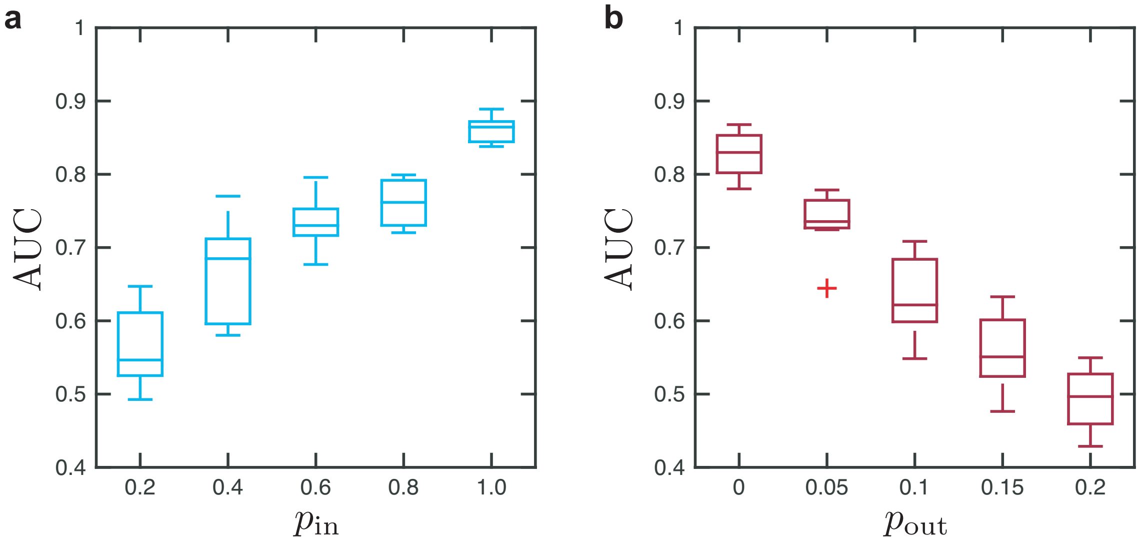

FIG. S10: **Performance of our DGM-based link prediction on synthetic networks generated by stochastic block models with varying within-cluster connection probability $p_{\text{in}}$ or between-cluster connection probability $p_{\text{out}}$.** The networks consist of 3 clusters/communities and each cluster has 16 nodes. Boxplots of AUC are calculated from 10 different realizations. For each realization, we randomly divide the links into two parts: a fraction of $10\%$ links chosen as the probe set and the remaining $90\%$ links as the training set. **a,** AUC as a function of $p_{\text{in}}$, with fixed $p_{\text{out}} = 0.05$. **b,** AUC as a function of $p_{\text{out}}$, with fixed $p_{\text{in}} = 0.5$.

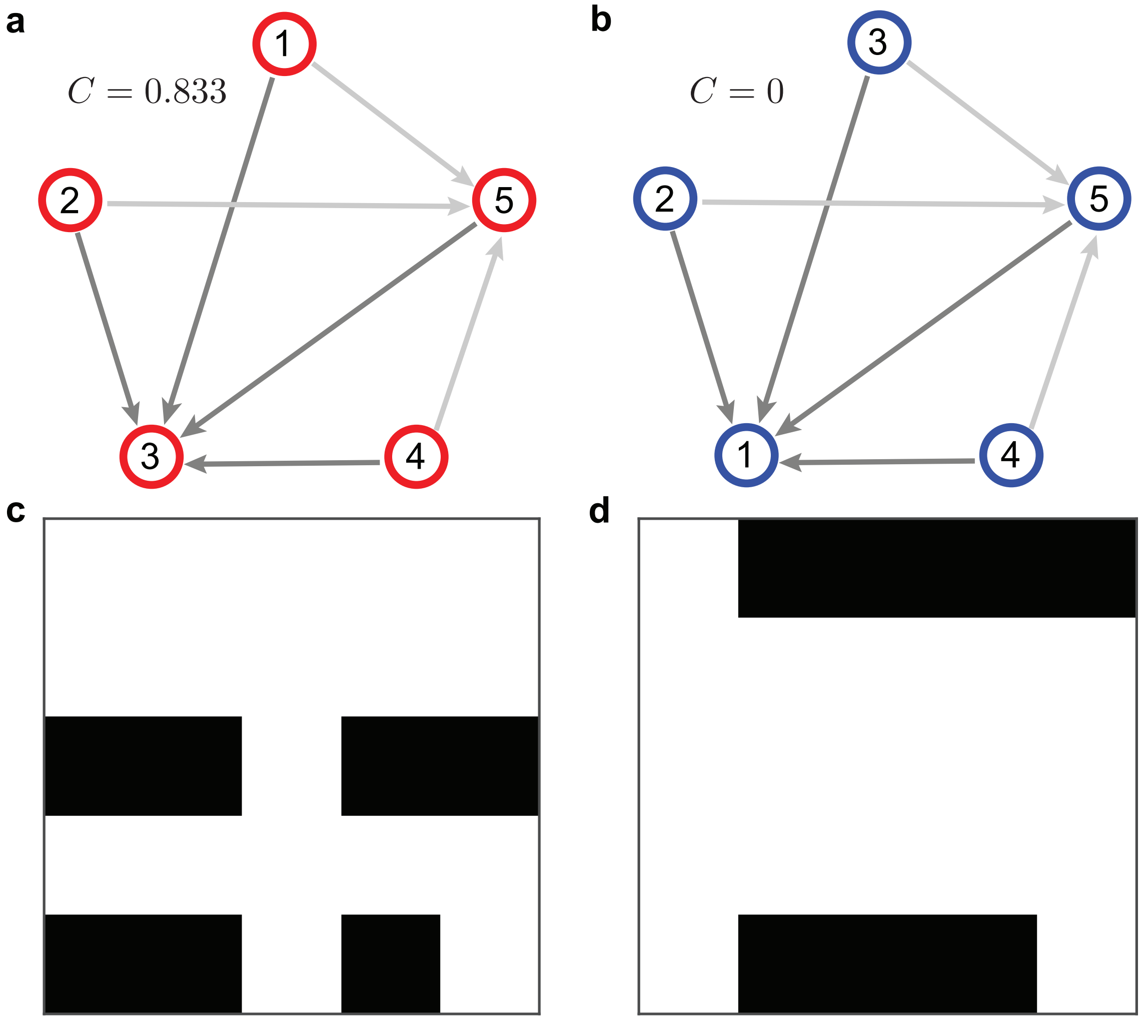

FIG. S11: **Illustration of our consensus-based node relabeling method.** To leverage any existing structural features in the network, we define a metric $E$: $E \equiv \sum_{i=1}^{N} \frac{\sum_{\alpha=1}^{k_i-1} |\Gamma(i)_{\alpha+1} - \Gamma(i)_\alpha - 1|}{k_i - 1}$, s.t. $k_i > 1$, where $\Gamma(i)$ is the neighbor set of node $i$ and $k_i = |\Gamma(i)|$. **a,** A directed network with 5 nodes and 7 links. The corresponding adjacency matrix is shown in (**c**). The metric $E$ for this network is $E = \frac{(2-1-1)+(4-2-1)+(5-4-1)}{3} + \frac{(2-1-1)+(4-2-1)}{2} \approx 0.883$. If we swap the labels of nodes 1 and 3, then the corresponding adjacency matrix is shown in (**d**). Then the matric $E = \frac{(3-2-1)+(4-3-1)+(5-4-1)}{3} + \frac{(3-2-1)+(4-3-1)}{2} \approx 0$. The global minimum $E^*$ is achieved and the network is shown in (**b**).

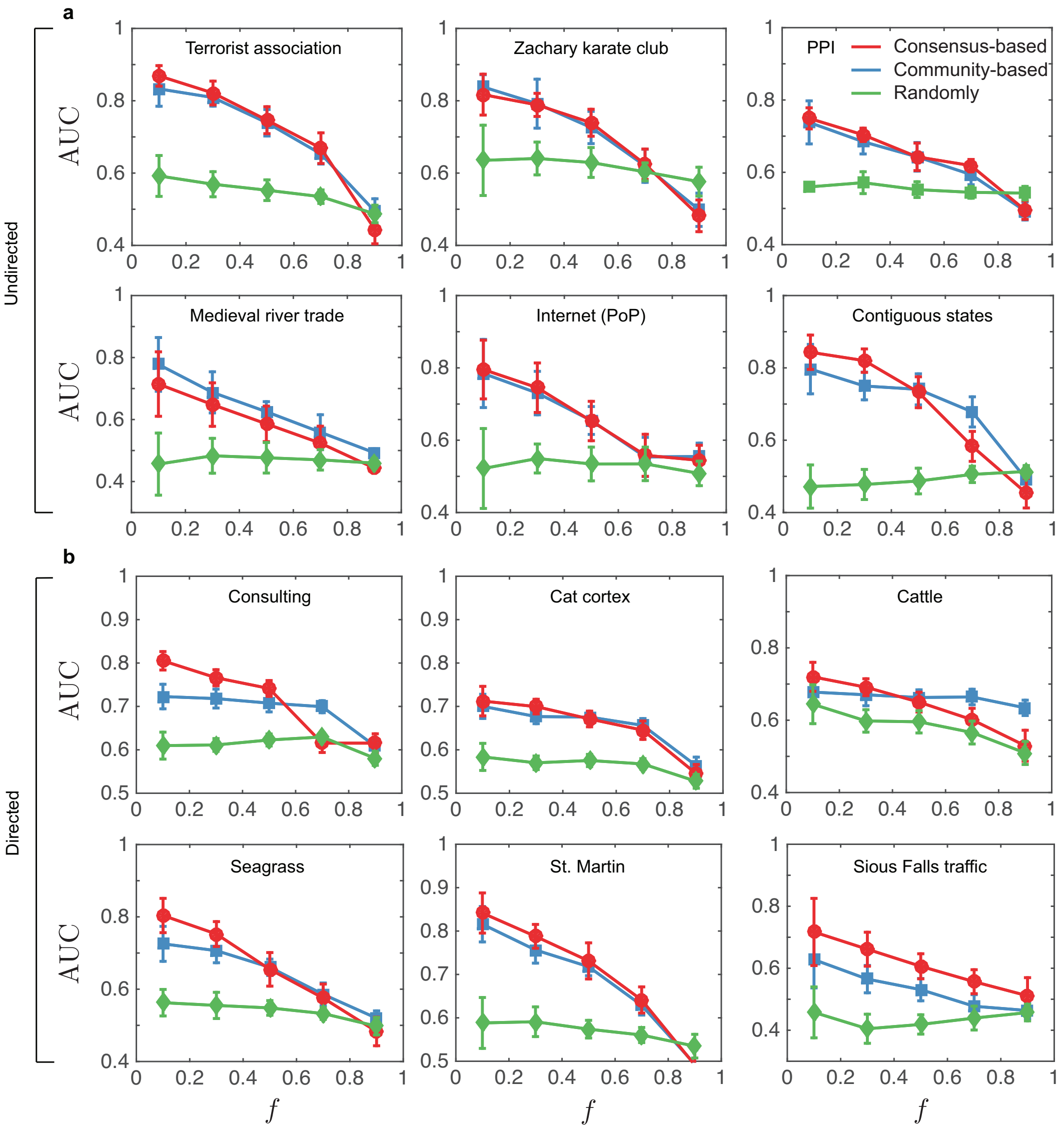

FIG. S12: **DGM-based link prediction with consensus-based node relabeling performs better than that with community detection-based node relabeling.** We compare the AUC of our consensus-based node relabeling (in red), community detection-based node relabeling (blue) and random node relabeling (green), respectively. Generally, our consensus-based node relabeling yields the highest AUC, especially for directed networks. **a,** Undirected networks. Top: Terrorist association networks, Zachary karate club, Protein-protein interaction network (a subnetwork of protein-protein interactions in *S. cerevisiae*). Bottom: Medieval Russian trade network, Internet (PoP level), Contiguous states (USA). **b,** Directed networks. Top: Consulting, cat cortex, cattle. Bottom: Seagrass food web, St. Martin food web, Sioux Falls traffic road. AUC of our DGM-based method is the average AUC over the last 20 epochs of the total epochs for all of the networks. Here we use GANs for the DGM.

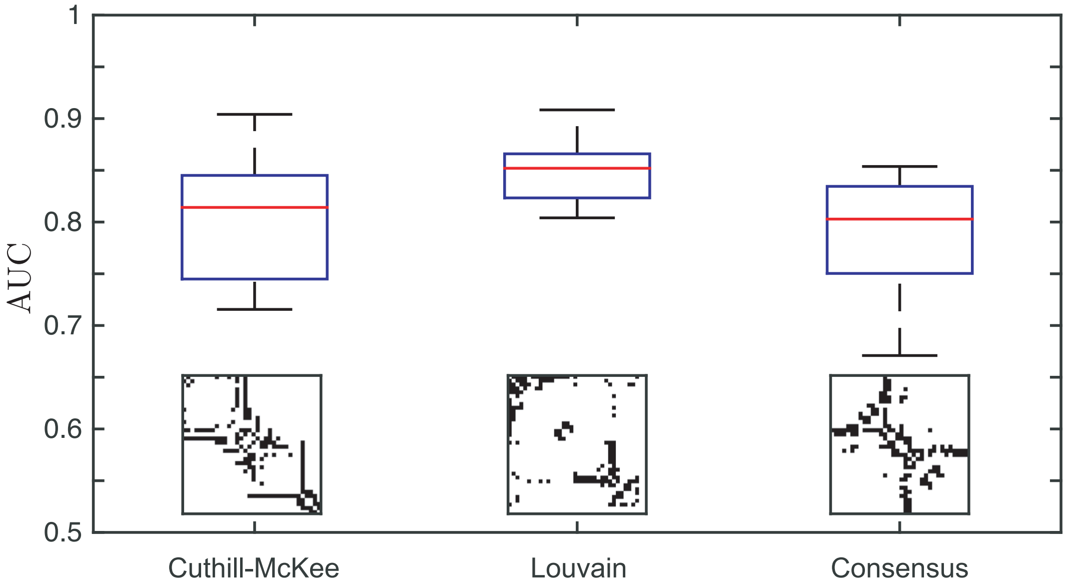

FIG. S13: **Performance of our DGM-based link prediction on an example network (Zachary karate club) labeled by three different methods: Cuthill-McKee algorithm, Louvain method, and our own consensus-based method.** Insets show the adjacency matrices reordered by the three methods. Boxplots of AUC are calculated from 10 different realizations. For each realization, we randomly divide the links into two parts: a fraction of 10% links chosen as the probe set and the remaining 90% links as the training set.

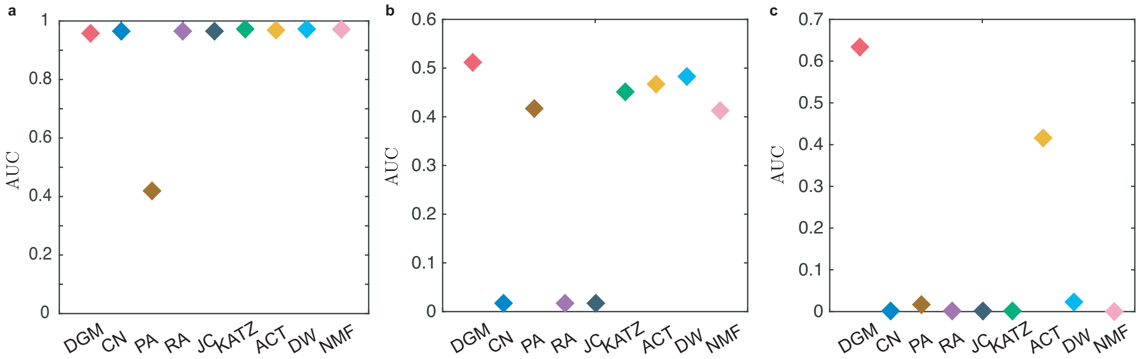

FIG. S14: **Performance of DGM-based link prediction and other scalable link prediction methods on modeled networks. a**, a synthetic stochastic block model network with 1056 nodes (with 12 clusters and 88 nodes for each, the connection probability of within cluster is $p_{\text{in}} = 0.5$ and between clusters is $p_{\text{out}} = 0.05$). **b,** random graph with connection probability $p = 0.01$ and **c,** random graph with connection probability $p = 0.001$. The network size is $N = 440$. We randomly divide the links into two parts: a fraction of 10% links chosen as the probe set and the remaining 90% links as the training set.

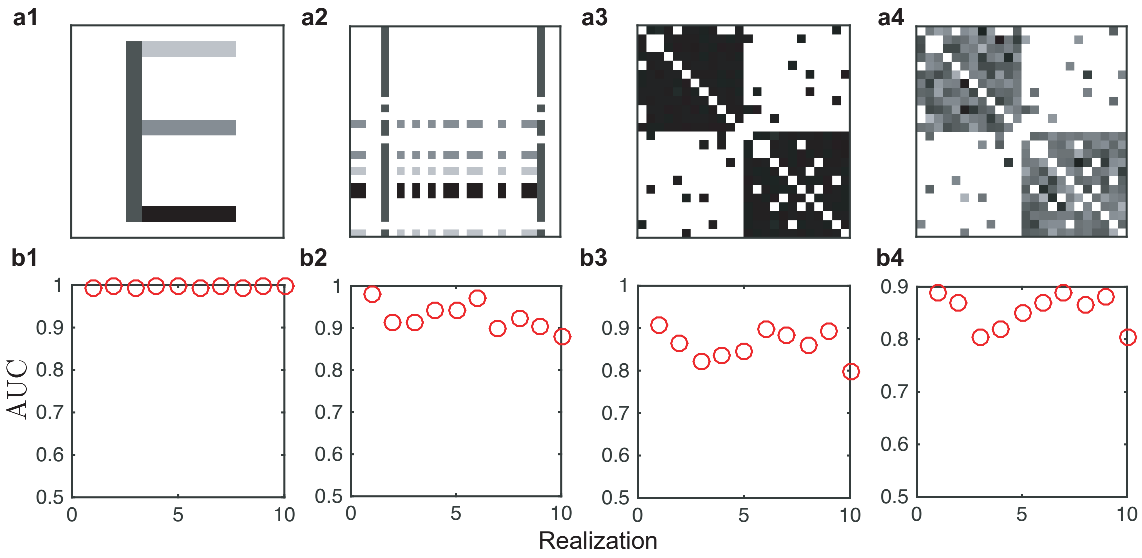

FIG. S15: **DGM in link prediction of weighted networks. a1,** The toy model of letter **E** and there have four link weights: 0.25, 0.5, 0.75, 1.0. **a2,** The network by randomly relabeling the node in (a1). **a3,** A stochastic block model with link weights drawn from normal distribution with mean 0.5 and derivation 0.01. **a4,** A stochastic block model with link weights drawn from normal distribution with mean 0.5 and derivation 0.1. **b1-b4,** AUC of DGM in 10 different realization for the networks in (a1)-(a4). We randomly divide the links into two parts: a fraction of $10\%$ links chosen as the probe set and the remaining $90\%$ links as the training set.

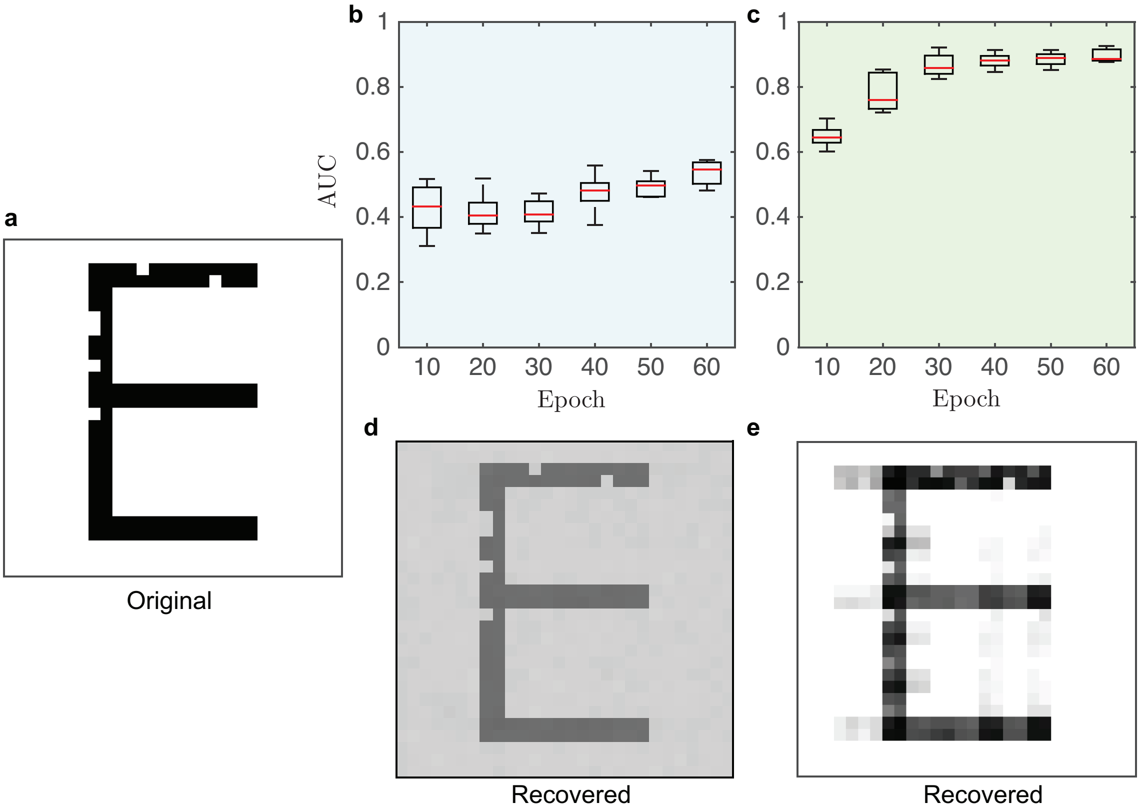

FIG. S16: **The VAE-based based link prediction generally does not perform as well as the GANs-based link prediction.** As in Fig. 1 of the main text, we are testing the link prediction for a directed network whose adjacency matrix has the shape of letter **E** (with some missing pixels). We divide the links of this network (with 28 nodes and 118 links) into two parts: $5\%$ be the probe set and the remaining $95\%$ be the training set. We find that the AUC of VAE-based link prediction (**b**) is considerably lower than that of GANs-based link prediction (**c**). **a,** Original network. **b,** AUC of VAE-based link prediction as a function of epoch. **c,** AUC of GANs-based link prediction as a function of epoch. **d,** Recovered network by VAE-LP. **e,** Recovered network by GANs-based link prediction.
